# Distribution-modeling quantifies collective Th cell decision circuits in chronic inflammation

**DOI:** 10.1101/2023.01.29.526090

**Authors:** Philipp Burt, Kevin Thurley

**Affiliations:** Systems Biology of Inflammation, German Rheumatism Research Center (DRFZ), a Leibniz Institute, Berlin, Germany; Institute for Theoretical Biophysics, Humboldt University, Berlin, Germany; Biomathematics Division, Institute of Experimental Oncology, University Hospital Bonn, Bonn, Germany

## Abstract

Immune responses are tightly regulated by a diverse set of interacting immune cell populations. Alongside decision-making processes such as differentiation into specific effector cell types, immune cells initiate proliferation at the beginning of an inflammation, forming two layers of complexity. Here, we developed a general mathematical framework for the data-driven analysis of collective immune-cell dynamics. We identified qualitative and quantitative properties of generic network motifs, and we specified differentiation dynamics by analysis of kinetic transcriptome data. Further, we derived a specific, data-driven mathematical model for Th1 vs. Tfh cell fate-decision dynamics in acute and chronic LCMV infections in mice. The model recapitulates important dynamical properties without model fitting, and solely by employing measured response-time distributions. Model simulations predict different windows of opportunity for perturbation in acute and chronic infection scenarios, with potential implications for optimization of targeted immunotherapy.

## Introduction

Immune responses are fast, highly dynamic and adapted to the specific pathogenic situation. Thereby, immune cells undergo collective decision-making processes, shaping the immune environment across conditions such as acute and chronic inflammation or the tumour microenvironment (1–5). For instance, peripheral T follicular helper (Tfh) cells drive B cell responses in synovial tissue derived from *Rheumatoid Arthritis* patients (2), and signaling between dendritic cells and natural killer cells is an important regulator of responsiveness toward anti-PD-1 therapy in human melanoma (6). Apart from direct cell-contact based interactions and more recently described mechanisms such as microvesicles, immune cells interact primarily via diffusible cytokines (7–12). Cytokines are proteins with diverse structures and biophysical properties, they often show pleiotropic functions, and they constitute a complex signaling network amongst immune cell populations.

Previous research has demonstrated the utility of mathematical models for gaining a functional, quantitative understanding of complex biological network dynamics. Well-studied examples come from metabolic, signal transduction and gene-regulatory networks (13–15). They have demonstrated that the functional properties of even simple network structures such as linear chains, negative or positive feedback and feed-forward loops, confound intuition and can only be understood by mathematical analysis and in relation to specific parameter values. Based on such insights, mathematical network models have for instance contributed to the successful development of targeted combination therapy strategies for perturbation of signaling pathways in cancer cells (16–19).

Despite the growing body of knowledge regarding pathway interactions and many recent breakthroughs in single-cell technology (20) and analysis methods (21–23), it remains a challenging task to perform quantitative studies on immune-cell decision circuits. On the one hand, acquiring kinetic data on all relevant system components is very difficult in situations where satisfactory in vitro systems are often not available. On the other hand, the very nature of the complex multi-scale problem spanning from intracellular signal transduction pathways to cell-cell interaction dynamics exacerbates convenient and easy-to-use mathematical formulations. In this context, we and others found that intracellular pathways are often well described using cellular input-to-output data in terms of measurable response-time distributions, instead of explicit descriptions of intracellular mechanisms (24–29). However, given the diversity of relevant processes, it remains unclear whether that approach is suitable to derive interpretable, data-based model formulations of a complex immunological process such as chronic inflammation.

Here, we designed a unified framework for data-driven mathematical descriptions of immune-cell population dynamics based on response-time modelling, capturing cell-cell interaction, cell proliferation and cell differentiation. Using that framework, we studied the generic behaviour of network motifs, we derived the structure of response-time distributions from a set of time-course transcriptomics data sets, and we performed detailed analyses of different strategies for population-growth control. Finally, we developed a specific data-driven model capturing the interactive Th cell dynamics observed during acute and chronic LCMV infection *in vivo*, and we employed that model to determine the relative effect of individual processes on the dynamics and to evaluate windows of opportunity for therapeutic intervention.

## Results

### A novel mathematical framework for modeling cell-population dynamics and decision-making reveals generic properties of immune-cell interaction circuits

Immune responses emerge from a collective process that is shaped by immune-cell communication, proliferation and differentiation. To mathematically analyze cell-cell communication processes, we previously found that cellular input-to-output dynamics are well described by response-time distributions quantifying the probability for individual cells to show a measurable response at time *t* to a stimulation at time 0 (24). Further, those response-time distributions can often be approximated using the two-parametric Erlang distribution (24,25,28). A process with such Erlang-distributed response times corresponds to an irreversible multi-step process, and vice versa, traditional rate-equation models on the level of cells correspond to exponentially distributed response times (30) (Figure S1A and B). Therefore, we will refer to that latter case as single-step models (SSM), and to the multi-step case as response-time models (RTM)(Figure S1C). Here, we generalized the previously developed formalism (24) to also cover cell proliferation and cell differentiation, or branching, into diverse cellular subtypes (Figure 1A and Figure S1A and Methods). Of note, we chose to explicitly set the probability for each branch in a decision tree as a function of the cytokine environment rather than as competition amongst response-time distributions (Figure S1D-G), since we reasoned that differentiation decisions can occur independently of response times. Overall, we designed a framework (Figure 1A) in which a cell can take a finite number of cell states, each with prescribed response-time distributions and probabilities for cell division and differentiation, which in turn are dependent on cytokine concentrations.

**Figure 1:**
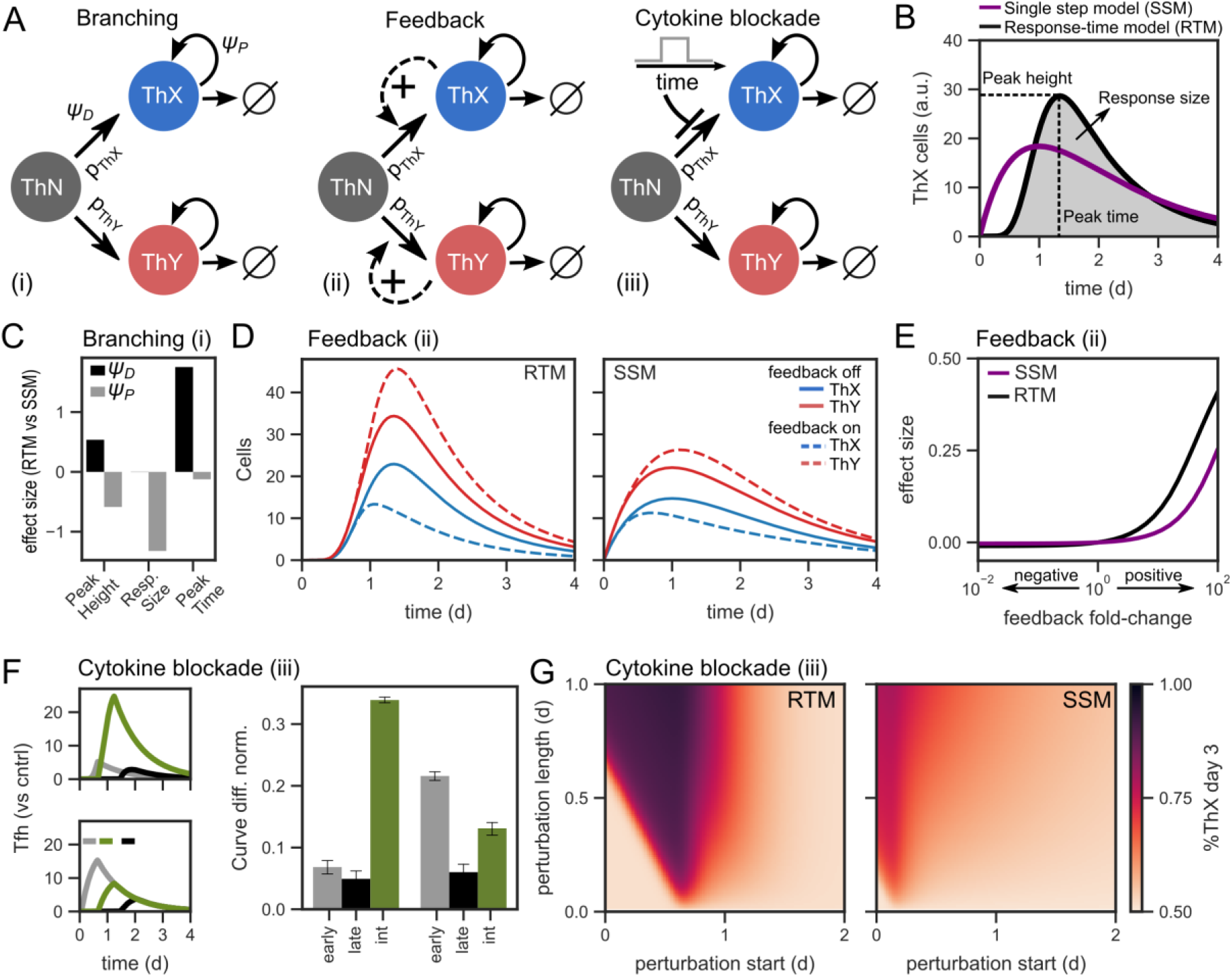
Properties of immune-cell interaction and proliferation circuits. (**A**) Generic models for branched cell fate-decisions and proliferation. The core motif consists of a naïve Th (ThN) cell population that differentiates into the proliferating effector sub-types ThX and ThY (see Supplementary Text S1 for details). (**B**) Time-course simulations resulting from the model shown in panel A (i). Inter-event-times between state-transitions are assumed to be either exponential (Single-step model; SSM) or Erlang-distributed (Response-time model; RTM), leading to different onset dynamics and quantifiable model readouts. (**C**) Effect on model readouts for the transition from an SSM model formulation (*k* = 1) to an RTM model (*k* = 10), the y-axis represents log2 fold-change of RTM vs. SSM dynamics. (**D-E**) Model branching dynamics with feedback (fold-change formulation, see Methods; cf. panel A (ii)). Shown are (D) model simulations with and without feedback on the branching probability, and (E) the log2 fold-change (effect size) in the ThX-to-ThY ratio as a function of the feedback fold-change. (**F**) Model dynamics exposed to transient perturbation, cf. panel A (iii). Left: Timecourse simulation of a cytokine blocking antibody released at time ***t_0_*** induces a change in the branching probability ***p_Thx_*** for a defined period Δt (time window indicated by colored patches above simulation). Shown are values normalized to a simulation without perturbation. Right: Change in total differentiated ThX cells for the same perturbation times as in the time-course simulations (color code in left panel). Error bars represent averages +/- SEM for simulations with randomized branching probability ***p_Thx_*** (n=50 simulations). (**G**) Systematic variation of perturbation length and start times. Change in ThX cells compared to control is indicated by color.

To explore how accounting for non-exponential state-transitions would affect response dynamics during cell proliferation and differentiation, we derived a binary fate-decision scenario, which can for example be interpreted as naïve T helper (Th) cells differentiating into the effector sub-types Th1 and Tfh cells, here labeled ThX and ThY (Figure 1A, model *i*). This simple branching model illustrates the key difference between the RTM and SSM models: multi-step dynamics as in the RTM formulation induce a delayed onset of the reaction and a sharp peak, while SSM formulations induce an immediate response of at least a small fraction of cells (Figure 1B). Based on these time-course simulations, we quantified the dynamics in terms of the properties ‘peak time’, ‘peak height’ and ‘response size’, defined as the area under the curve (Figure 1B). The response size quantifies the total number of cells generated during the immune response under study, while the other properties characterize the speed and the maximal intensity of the response of the analysed immune cell population. As a result, when changing from an SSM to an RTM model formulation by increasing the step number *k* for differentiation, we observed increased peak times and peak heights (cf. Figure 1B), whereas the overall response size was not affected (Figure 1C). In contrast, increasing *k* in the response-time distribution describing cell proliferation did not only lead to changes of the peak time and height, but also induced pronounced effects on the response size (Figure 1C). Hence, a traditional model formulation using rate-equations would overestimate the resulting Th cell population size.

Next, to analyze cytokine-mediated positive feedback, which has been described for instance in the case of IFN-gamma mediated differentiation into Th1 cells (31), we assumed a symmetric feedback scenario where each effector cell type secreted a cytokine species with positive feedback on the respective branching probability *p_Thn,ThX_* or *p_Thn,ThY_* (Figure 1A, model ii). In both the RTM and SSM model variants, the cytokine-induced feedback amplified initial differences in the ThX-to-ThY ratio, although differences were less pronounced in the SSM model (Figure 1D and E). Apart from feedback induced by already differentiated cells, also cytokine signals derived from other cell types, or cytokines or cytokine blockers that are present in terms of therapeutic interventions, may affect cell-fate decisions. To investigate that scenario, we considered a time-dependent modulation of the branching probability (Figure 1A, model iii). We proceeded to evaluate the perturbation regarding effectiveness of dynamical treatment conditions such as early vs. late starting times after stimulation, and we found that optimal perturbation times varied between the RTM and SSM models. In particular, while early intervention was most effective in the SSM scenario, the RTM model revealed distinct optimal perturbation windows at intermediate perturbation times (Figure 1F and G). Late perturbation times resulted in small effect sizes compared to both early and intermediate perturbation times in both the SSM and RTM models. The effectiveness of intermediate perturbation times in the RTM model was largely independent of perturbation-specific parameters such as the initial difference in branching probability (error bars in Figure 1E, right panel), as well as model-specific parameters such as the shape parameter of the corresponding response-time distribution (Figure S1H-I).

Overall, we found that cellular decision-making and proliferation dynamics depend sensitively on the shape and scale of the input response-time distribution, thus emphasizing the need to account for details of those distributions in quantitative models of specific systems.

### Measured gene-expression kinetics in Th cell decision-making are well described by gamma or exponential distributions

We next asked to what extent measured response-time distributions for Th cell dynamics are well described by our framework. To this end, we performed a systematic analysis of the kinetic properties of differentiation dynamics across several kinetic transcriptome data sets (Figure 2A, Methods). In particular, we derived the best-fit gamma distribution model *γ*(*α,β*) to the normalized kinetic genes in each data set and classified a gene as gamma-distributed if the fit was significantly better compared to the fitting statistics derived from an exponential distribution model (Figure 2B, Figure S2A-E, Table S1). Of note, the fitting quality did not show substantial variation when using a less detailed description by an Erlang distribution with integer-valued shape parameter, as used in our modelling framework, instead of the corresponding best-fit gamma-distribution (Figure S2F). Genes with poor fitting statistics were further grouped into the category “other”, or into the category “bimodal” if instead they showed good fitting statistics to a mixed Gaussian model (Figure S2G). We found at least 70% of the kinetic Th cell genes to be well-described by either gamma or exponential distributions across all studied relevant gene-sets, and ~50% of those genes were significantly better described by choosing a non-exponential gamma distribution with *α* > 1 (Figure 2C and 2D). Of note, we detected significant enrichment for cell-cycle signaling and metabolism predominantly in the “gamma” category (*α*>1) across all data sets, highlighting the functional contribution of those genes (Table S2).

**Figure 2:**
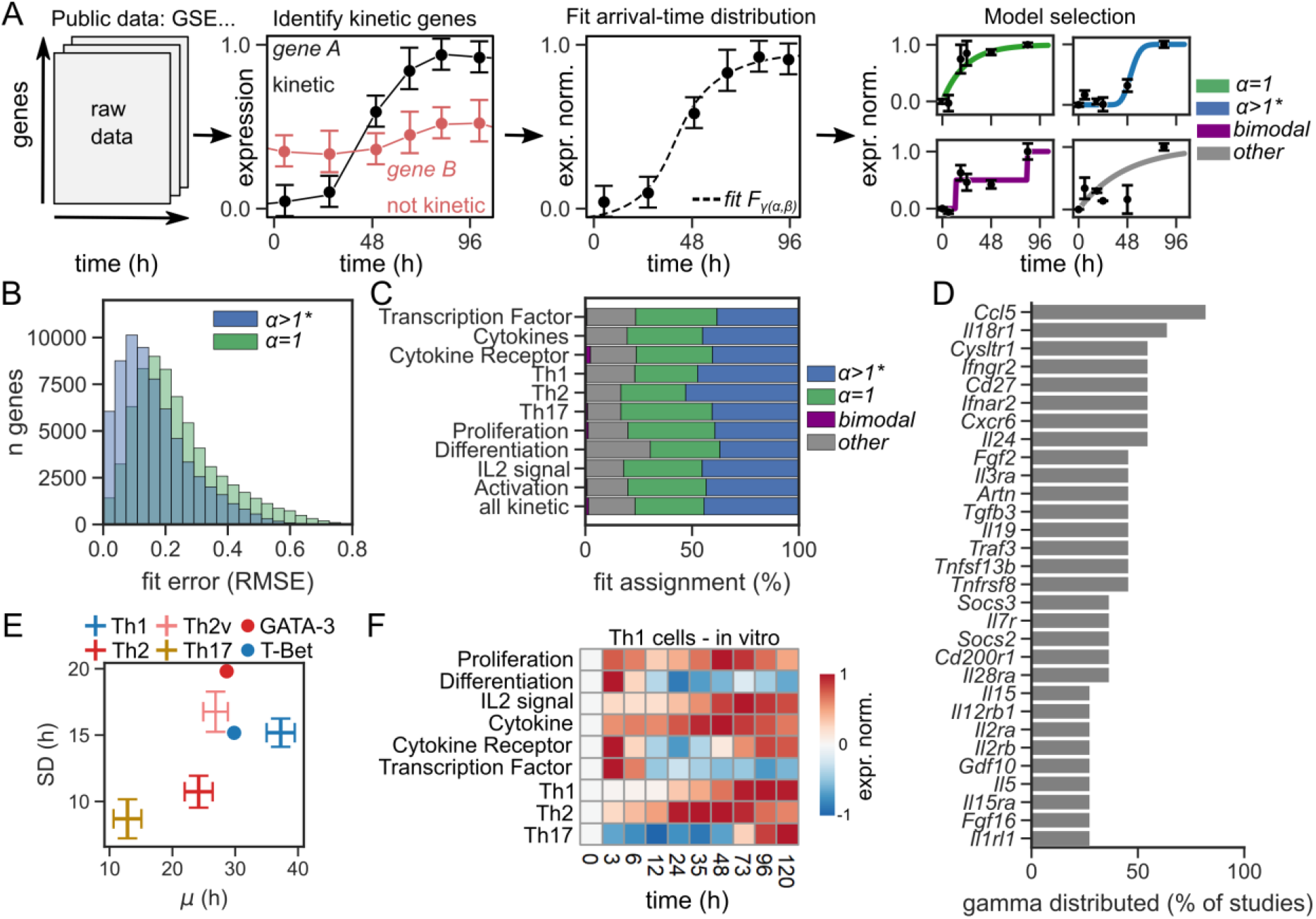
Quantitative analysis of Th cell time-course transcriptomes. (**A**) Analysis workflow. Kinetic gene expression data from public repositories (Table 2) are filtered for genes with significant expression changes over time. Those kinetic genes are then normalized and fitted to an arrival-time distribution *F*_*γ*(*α,β*)_, and kinetic genes are assigned to categories based on the best-fitting kinetic model (data re-plotted from (60)) (see Methods for details). (**B**) Distribution of the root-mean-square error (RMSE) of fitting procedures across kinetic genes in all analyzed data sets for SSM and RTM models. (**C**) Category assignment for different gene modules resulting from the described work-flow. (**D**) Top 40 genes assigned as gamma-distributed for the gene modules shown in panel D. (**E**) Average *μ* and standard deviation SD for best-fit parameters *α*, *β* of individual effector cell differentiation modules (see Methods). (**F**) Normalized expression values for gene modules in Th1 cells.

Analysis of individual Th cell signature genes across data sets (Figure 2D) indicated a high percentage of delayed kinetics (*α*>1), especially for certain cytokine and chemokine receptors. Further, based on the derived gamma-distribution fitting parameters, we aimed to categorize and quantify Th cell relevant kinetic genes in more detail. Focusing on a set of specialized Th cell gene lists (32), we were able to identify average transition times and standard deviations for each pathway by averaging the parameter estimates over all kinetic genes within a specific gene set (Figure 2E and Figure S2H). Interestingly, the gene modules showed quite specific kinetics ranging from stable up- or downregulation to transient expression dynamics (Figure 2F, Figure S2I).

Taken together, our analysis indicated that the majority of kinetic genes, including highly important genes for Th cell differentiation and communication, are well described by gamma distributions with onset kinetics that are quite specific for individual functional gene modules.

### Quorum sensing and timer mechanisms are required for stable proliferation dynamics

To arrive at a specific data-driven formulation of immune cell dynamics in the context of inflammation onset, we next studied physiological mechanisms for the regulation of cell proliferation. It is well-established that cellular turnover rates need to be tightly regulated by feedback mechanisms, since a system with independent birth and death rates is structurally unstable and inevitably causes either extinction or unbounded cell growth (Figure 3A, “naive”) (33). Thus, we asked whether physiologically plausible mechanisms could ensure stable expansion in the context of varying proliferative capacities and under conditions with re-stimulation. For this purpose, we assumed that activated naive cells transition into precursor cells and finally into effector cells, and we took the division rate as a control parameter that we assumed to be regulated by the activation level of the cell population. We refer to Th cell dynamics here, but other proliferating immune-cell populations such as CD8+ T cells (34,35) and NK cells (36) have been described by similar mathematical formalisms.

**Figure 3:**
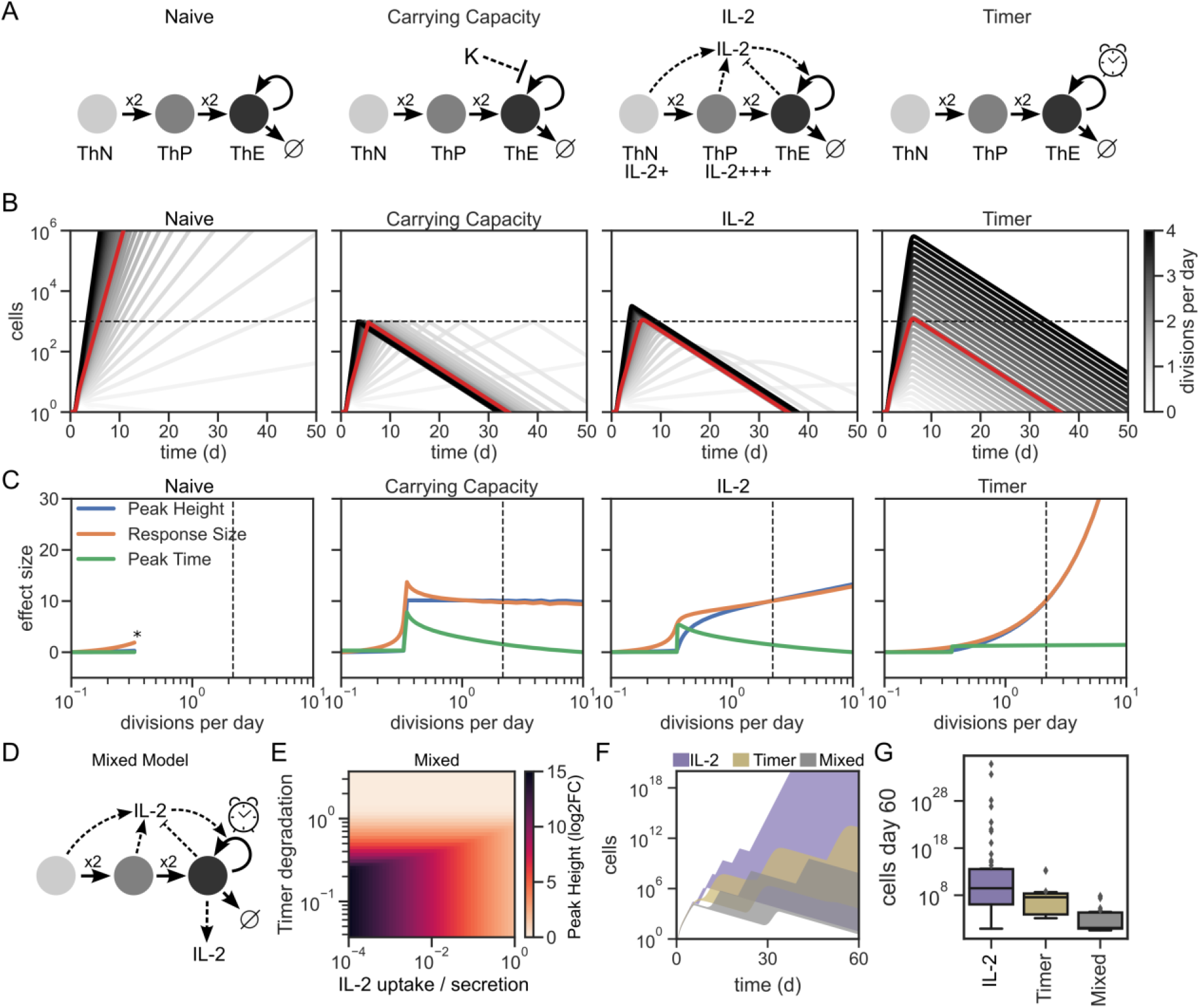
Mechanistic models of cell-population growth control. (**A**) Plausible model designs for cell-population growth control. (**B**) Effector Th cell dynamics based on model simulations for the models shown in (A) for different values of the proliferation rate. Model-specific-parameters were normalized to the same response magnitude (red line). (**C**) Quantification of peak height, peak time and response magnitude of the simulations shown in (B) (cf. Figure 1B for an illustration of model readouts). Values represent log2-fold-changes with respect to a model simulation with default parameter values. * indicates lack of stability, i.e. monotonically increasing cell numbers for at least 100 days. (**D**) Combination of the IL-2 and Timer models (panel A) into a ‘Mixed Model’. (**E**) Sensitivity of the response peak height in the Mixed Model for variation of model-specific parameters (x-axis: IL-2 model, y-axis: Timer model). Peak height displayed as log2-fold-change compared to the minimal peak height. (**F**) Model simulations with restimulation in the IL-2, Timer and Mixed Models. Filled areas represent the maximum range of observed dynamics. (**G**) Distribution of cell numbers at day 60 for the simulations shown in (F).

We compared three different minimal models of regulated effector proliferation to the ‘naïve’ scenario (Figure 3A): (i) requirement for uptake of external resources (‘carrying capacity’), (ii) limited availability of a cytokine such as IL-2, which is secreted by naïve and precursor cells themselves (‘IL-2 model’), or (iii) a constraint on the absolute time passed since Th cell activation (‘Timer model’) (33,37,38). Of note, a mechanism that depends on the size of one or several of the involved cellular compartments, such as the IL-2 model, has often been termed a quorum sensing mechanism in the literature (39). Analyzing the kinetic properties of a cell population (cf. Figure 1C), we found that the ‘naive’ model generated unstable dynamics already for slight modulation of the division rate, as expected (Figure 3B and 3C). The ‘carrying capacity’ mechanism allowed for robust control of the population size, but it fell short regarding adaptability of the resulting dynamics to the specific needs of a concrete immune response as mimicked by modulation of the division rate. The ‘IL-2 model’ and the ‘Timer model’ both generated an appreciable dynamic range regarding control over peak height and response size for intermediate values of the division rate (Figure 3C). The ‘IL-2 model’ further allowed for simultaneous control over the peak time, while that property was fixed by construction of the ‘Timer model’. Further, the ‘Timer model’ showed somewhat higher plasticity of the response size, but also had a tendency to become unstable for very high division rates.

Hence, the ‘IL2 model’ appeared to outperform the other hypothesized mechanisms regarding robustness and tunability. However, we reasoned that a quorum sensing mechanism such as the ‘IL2 model’ alone might be prone to instability in the presence of repetitive stimulation as in the case of chronic inflammation, because in each cycle, some IL-2 secreted in a previous cycle can be expected to remain in the system. Indeed, our simulations supported such a scenario in a regime with high but still physiologically plausible IL-2 secretion rates of 150 molecules cell^-1^ s^-1^ (40) (Figure S3A-B). Therefore, we proceeded to analyse a combination of the ‘IL-2 model’ and the ‘Timer model’, and we found that the resulting ‘Mixed model’ exhibited robust buffering properties also in the case of repetitive stimulation (Figure 3D-G and Figure S3).

Thus, the combination of time-dependent and quorum-sensing mechanisms ensured stable expansion even in heterogeneous environments with different interclonal proliferative capacities and recurring antigen encounters.

### A data-driven model of Th cell decision-making in acute and chronic inflammation

A critical compartment of many immune cell decision-making processes are diverse lineages of Th cells, which can promote both cellular immune responses and antibody-mediated responses, depending on their phenotype(31). Even for very similar pathogenic scenarios such as closely related viral infections, the dominant effector Th cell phenotype can vary substantially. In acute viral infections, Th cell responses are typically dominated by Th1 cells, promoting a strong cellular immune response. In contrast, during chronic viral infections such as Hepatitis C virus (HCV) or human immunodeficiency viruses (HIV), the landscape of Th cell phenotypes is more diverse and often comprises expanded T follicular helper (Tfh) cells promoting the development of antibody-producing plasma cells (41). Additionally, Th cells undergo a phenotype alteration, exhibiting reduced effector functions such as loss of proliferation potential, reduced effector cytokine secretion, upregulation of inhibitory markers and secretion of the immunosuppressive cytokine IL-10 (42,43). Here, we focused on Lymphocytic choriomeningitis virus (LCMV) infection in mice, a well-established experimental model system for the study of acute and chronic viral infections (43–45).

To derive a quantitative model of Th cell responses in acute and chronic inflammation, we built on two considerations derived from recent experimental data. First, it has been shown that precursor cells maintain a level of reduced proliferative capacity which can sustain the development of fully differentiated cells, even during chronic inflammatory conditions (34,46). Second, terminally differentiated Th1 cells secrete the cytokine IL-10, constituting a feedback circuit modulating the frequency of Tfh cells (47). Based on those constraints, we designed a closed model formulation by extending the ‘Mixed model’ (cf. Figure 3) and additionally considering differentiation of naive Th cells into Th1 and Tfh effector cells as well as further development into chronic and memory cell states (Figure 4A). Although LCMV infection in mice has been studied experimentally in much detail, kinetic data especially for early time-points at the onset of inflammation is not widely available. For data annotation, we therefore initially referred to published data-sets with high-resolution *in vitro* kinetics to determine the response time distributions for cell division and differentiation, as well as the rate parameters for cellular half-lives and cytokine secretion rates (Figure 4B, Table 1 and Table S3) (8,40,48–50). We found that after this pre-conditioning step, a published *in vivo* data-set from LCMV-infected mice (51) with only 3 time-points per condition was sufficient to determine the remaining parameters up to a narrow range around the best-fit curve (Figure 4C-D). This annotation strategy strongly reduced the range of observed dynamics in the annotated model version, both compared to a model where all parameters were determined by best-fit estimation using the *in vivo* data set (RTM), and when compared to a model that assumed exponential inter-event-times (SSM) (Figure 4E-F). In particular, the reduced and stable levels of effector cells, a hallmark of chronic inflammation, were reflected by design of the model after annotation using only the response-time distributions derived from in vitro data (dark shaded regions in Figure 4E). Fitting of Th1 and Tfh cell numbers to a specific ex vivo data set merely served to further fine-tuning the dynamics and providing more robust inference of the remaining fitting parameters (Figure 4G).

**Figure 4:**
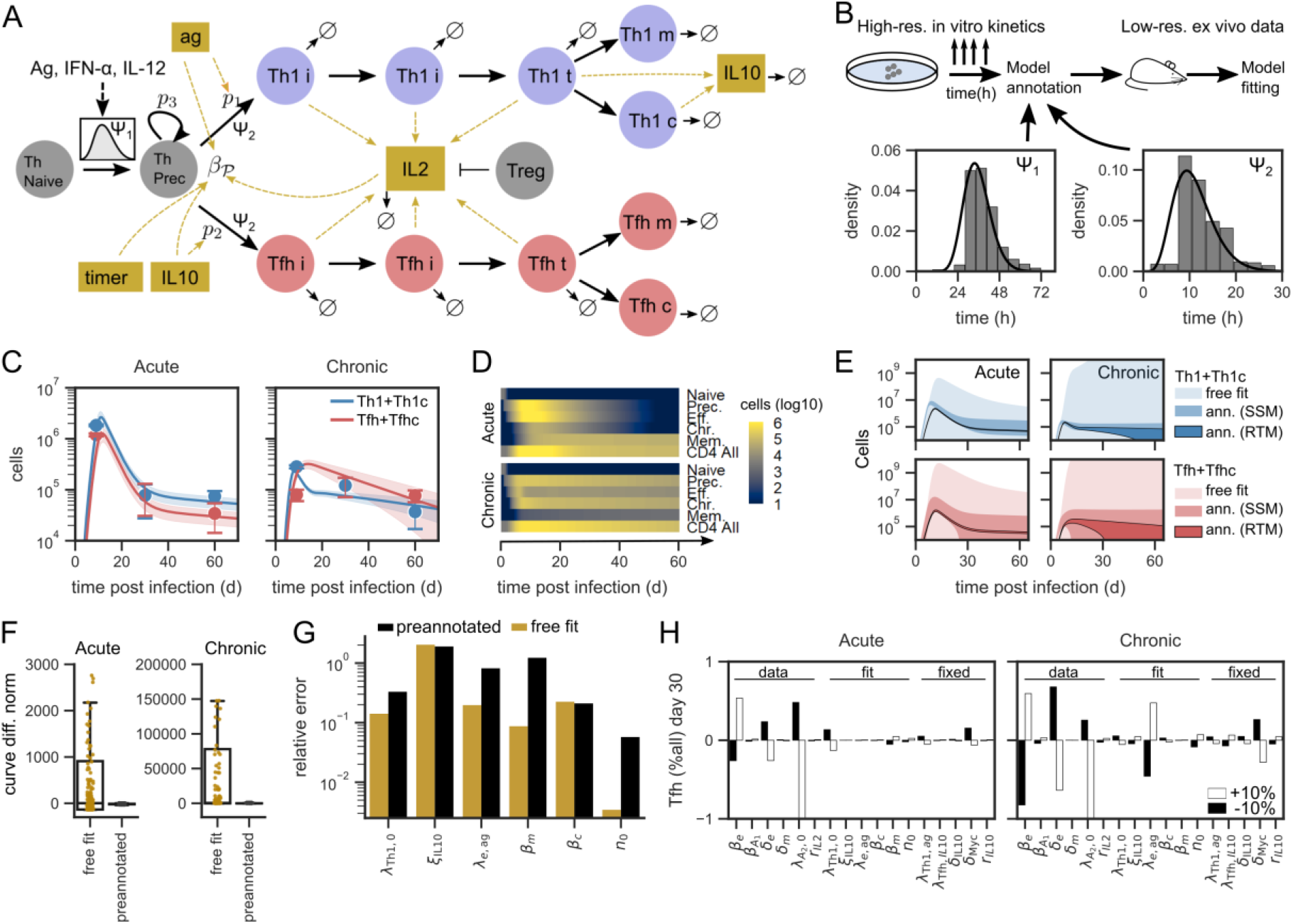
Specific response-time model of Th cell dynamics in acute and chronic LCMV infection in mice. (**A**) Model Scheme. Naive Th cells differentiate into proliferating precursor cells, which can differentiate into effector cells (Th1 and Tfh) or chronic cells (Th1c and Tfhc). Effector cells can further transition into memory cells. The probability to obtain a given cell-fate is modulated by concentrations of IL-10 and antigen. (**B**) Workflow for model annotation. High-resolution in vitro data is used to parameterize Erlang distributions describing proliferation and differentiation dynamics. The resulting reduced set of parameters is determined by curve-fitting to ex vivo data (Table 1). (**C**) Combined model fit to acute and chronic viral infection kinetics, shown are best-fit model dynamics and corresponding data for total Th1 and Tfh cells (data replotted from Ref. (51)). Shaded areas represent SD of model dynamics with model parameters being drawn from a lognormal distribution (CV=0.05, average set to bet-fit parameter value). (**D**) Dynamics of individual cell subsets for the best-fit model simulations in (C). (**E**) Range of model dynamics for Th1 and Tfh cells for indicated annotation strategies, shaded area as in (C) but with parameters drawn from a uniform distribution bounded by the **2σ–**confidence interval for the respective model fit. (**F**) Curve-difference between the best-fit simulation and the simulations with parameters drawn from random numbers shown in (E). (**G**) Relative error for individual model parameters for the free fit model and with the annotated model version (RTM model).

**Table 1:**
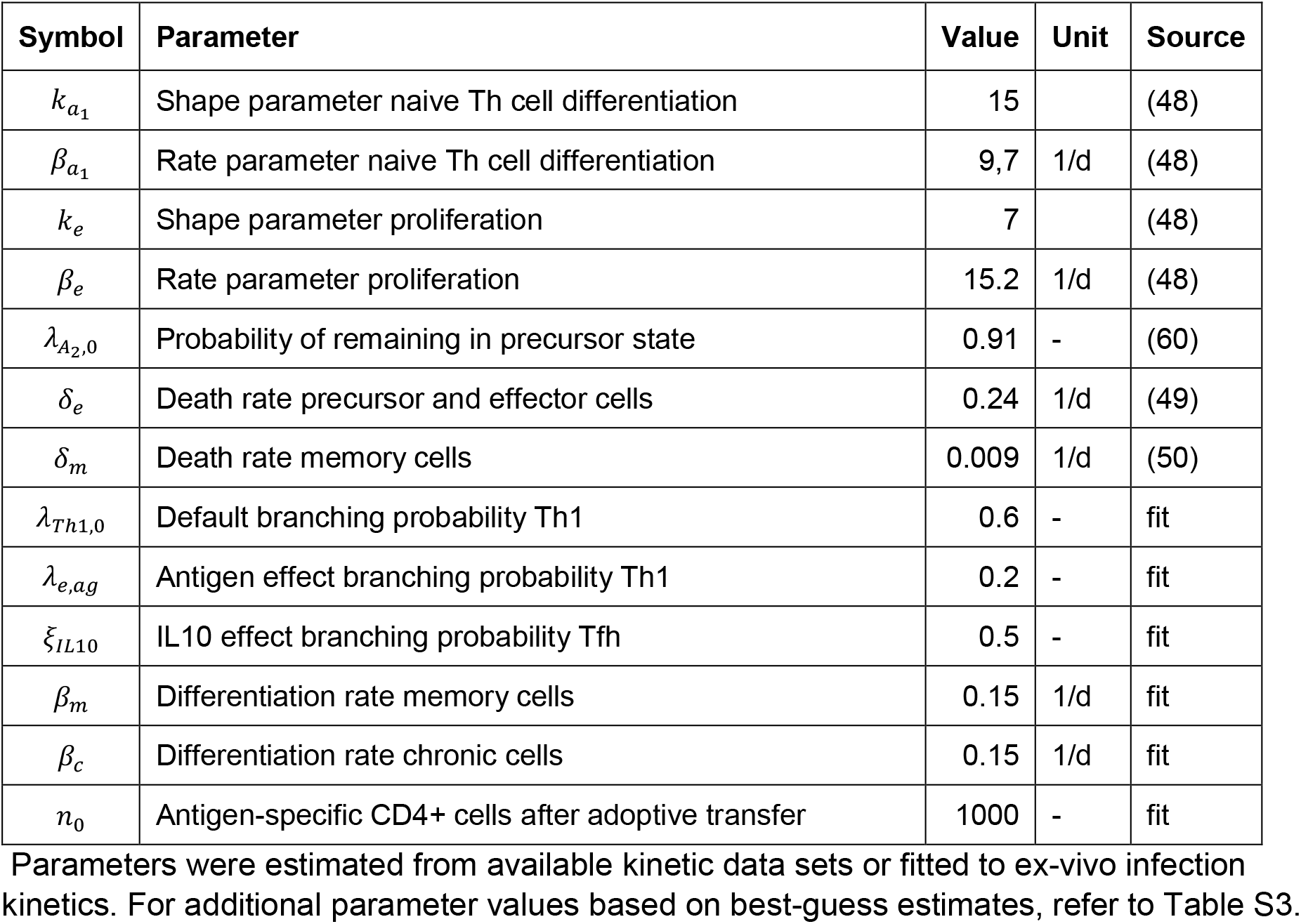
Parameter values used in the mathematical model.

| Symbol | Parameter | Value | Unit | Source |
| --- | --- | --- | --- | --- |
| $k_{a_1}$ | Shape parameter naive Th cell differentiation | 15 | | (48) |
| $\beta_{a_1}$ | Rate parameter naive Th cell differentiation | 9,7 | 1/d | (48) |
| $k_e$ | Shape parameter proliferation | 7 | | (48) |
| $\beta_e$ | Rate parameter proliferation | 15.2 | 1/d | (48) |
| $\lambda_{A_2,0}$ | Probability of remaining in precursor state | 0.91 | - | (60) |
| $\delta_e$ | Death rate precursor and effector cells | 0.24 | 1/d | (49) |
| $\delta_m$ | Death rate memory cells | 0.009 | 1/d | (50) |
| $\lambda_{Th1,0}$ | Default branching probability Th1 | 0.6 | - | fit |
| $\lambda_{e,ag}$ | Antigen effect branching probability Th1 | 0.2 | - | fit |
| $\xi_{IL10}$ | IL10 effect branching probability Tfh | 0.5 | - | fit |
| $\beta_m$ | Differentiation rate memory cells | 0.15 | 1/d | fit |
| $\beta_c$ | Differentiation rate chronic cells | 0.15 | 1/d | fit |
| $n_0$ | Antigen-specific CD4+ cells after adoptive transfer | 1000 | - | fit |

To further determine the effect of individual model parameters, we analyzed the frequency of Tfh cells on day 9 post activation (Figure S4A), where chronic infection is beginning to emerge, and on day 30, where chronic infection is fully established (Figure 4H). We found that during both acute and chronic infection, differences in the Th1-to-Tfh ratio were to a large extent driven by processes regulating cellular turnover such as proliferation and cell death, which in the model are described by response-time distributions obtained from in vitro data (Table 1). Parameters identified by fitting ex vivo data further specified the dynamics. Especially, variation of the branching probability coefficient *λ_Th1,0_* during acute infection and variation of the antigen coefficient *λ_e,ag_* during chronic infection had a substantial effect (Figure 4H and Figure S4B). Parameters without data annotation only showed a minor contribution except for regulation of the timer-intrinsic parameter, which had a strong effect at early time points (day 9 post infection) in acute infection but not in the chronic infection scenario.

Taken together, our mathematical framework allowed designing a data-driven model of Th cell interaction in acute and chronic inflammation which was to a large degree constrained by quantitative formulations of individual processes, and further refined by post-hoc data fitting.

### Quantitative predictions of relapse effects and effective intervention times

The data-annotated model allowed us to revisit the previously observed effects of transient signals (cf. Figure 1E-F), which may be present in the form of other interacting cell types, cytokines or cytokine blockers. To analyze such a timed perturbation in the specific scenario of acute and chronic LCMV infection in mice, we used the best-fit model parameterization and modulated the probability to adopt a Th1-cell fate within a given period of time (Figure 5A). During acute infection, the Th1-to-Tfh ratio was most susceptible to intermediate perturbation times and was irresponsive to late perturbation times, which is in agreement with our previous analyses using the minimal branching models. Interestingly, during chronic infection, perturbations also induced shifts in the Th1-to-Tfh ratio at later time points. However, after withdrawal of the stimulus, the Th1-to-Tfh ratio returned to its prior unperturbed value (Figure 5B, Figure S4B). Thus, the cells exhibited a “relapse” effect with short-lasting perturbation effects, which can be attributed to the antigen-dependent continuous proliferation of precursor cells. Long-lasting effects could only be achieved at very early perturbation times, but in that case the effect size was much smaller compared to all other scenarios under study (Figure 5B). We systematically investigated the effect of timed perturbations by varying the start time and duration of a perturbation in our simulations (Figure 5C and D, cf. Figure 1C), and we found that the relapse effect occurred largely independently of both parameters. In particular, combining both parameter scans gave rise to a tongue-like contour plot (Figure 5D) showing that strong changes in the Th1-to-Tfh ratio only occur during limited time-intervals as “Response maximum”, while in the acute inflammation scenario, also the average effect size over the whole simulated time-course is modulated.

**Figure 5:**
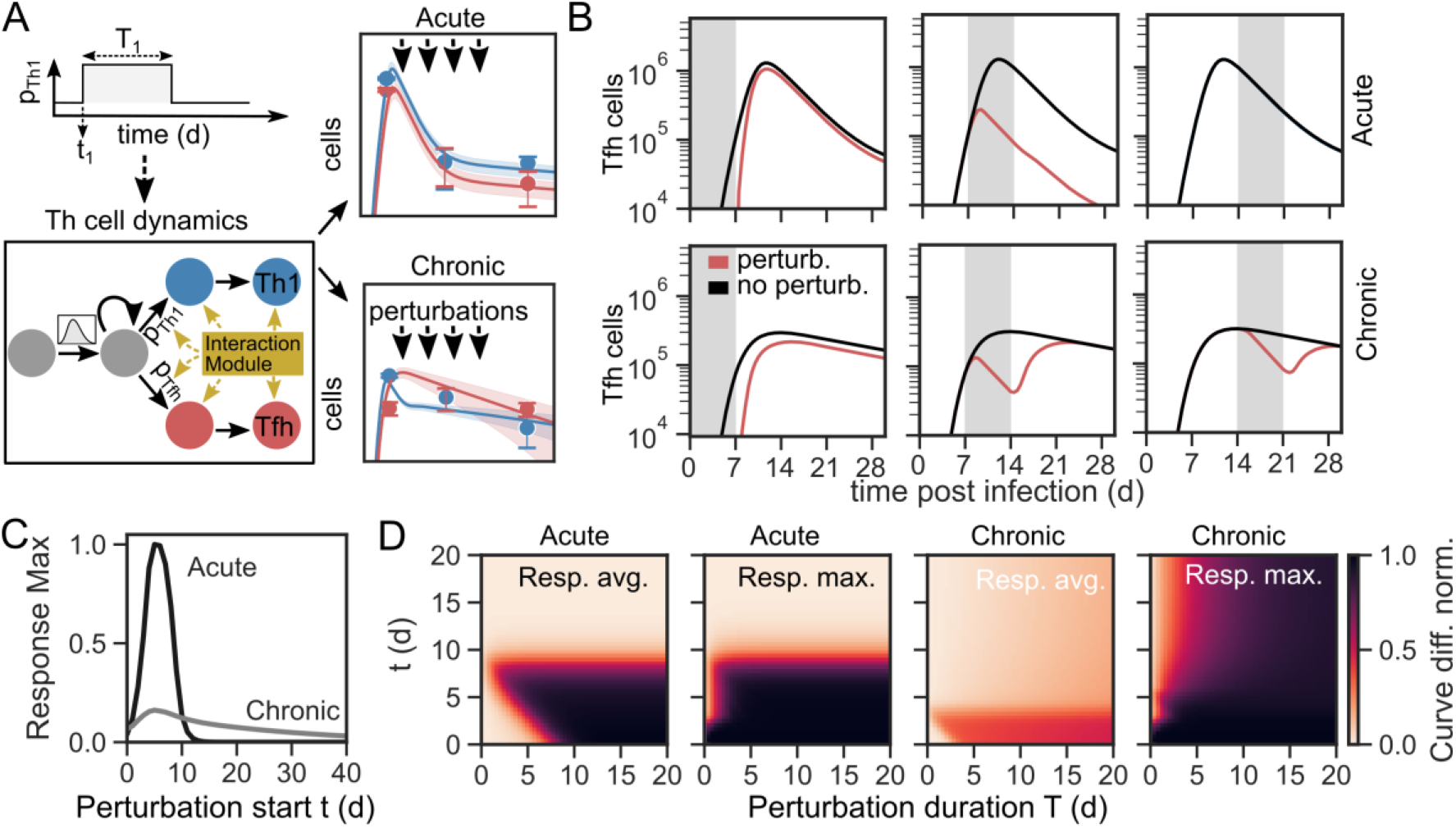
Analysis of optimal time-windows for perturbation of immune-cell dynamics. (**A**) Workflow to identify optimal perturbation times. Branching probabilities are varied within a time-window defined by the start time *t* and the duration *T* of the perturbation in the best-fit model (cf. Figure 4C and Supplementary Text). (**B**) Model simulations for early, intermediate and late perturbation times with a duration of *T*=7d for acute and chronic infection. Shown are the numbers of Tfh cells. (**C**) Effect of perturbation onset during acute and chronic infection. Shown is the maximum number of Tfh cells, normalized to a simulation without perturbation. (**D**) Systematic variation of perturbation length and start times during acute and chronic infection. Shown are the change in average levels (“Resp. avg.”) and maximal change (“Resp. max.”) in the number of Tfh cells over time compared to a simulation without perturbation (min-max normalized).

Overall, our model simulations predicted pronounced differences between acute and chronic inflammation scenarios for timed perturbation schemes in terms of both response size and dependency on the perturbation onset time.

## Discussion

In this study, we introduced a generic framework for the mathematical analysis of Th cell fate-decisions alongside proliferation and cell-cell communication using information from single-cell reaction dynamics. While the framework is tailored towards Th cells, it can be flexibly extended to encompass alternative cell types and fate-decisions. The adopted approach is based on ordinary differential equations instead of stochastic dynamics, so that numerical simulations can be performed with high performance, allowing for systematic analyses of many different parameter settings even in large and complex model implementations. We used the developed modelling framework to study a set of frequently occurring network motifs, to compare plausible mechanisms of cell-growth control at the onset of inflammation, and to design a data-driven model of Th cell dynamics in LCMV infection in mice.

Analysis of published time-course transcriptome studies revealed that our modelling framework captures the onset dynamics of the vast majority of kinetic genes. Around 50% of those genes were best described by a model with non-exponentially distributed inter-event times, clearly indicating a need for distribution-based modelling instead of traditional rate equation models for the analysis of cell-cell interaction dynamics. Notably, generalization of our approach to considering also multi-modal distributions as input data is straight-forward and merely requires a linear chain formalism where the assumption of identical rate parameters for all intermediate steps is relaxed (28,52). Our results are in line with previous studies showing that cellular reaction times are often non-exponential (24,25,53), and therefore are better described using non-Markovian dynamics, especially if detailed quantitative knowledge on intracellular processes is not available.

Immune-cell proliferation dynamics is a crucial process for maintaining functional immune responses and thus requires particular consideration when conceptualizing a quantitative formulation. Our analysis of response-time models of cell proliferation emphasized the previous finding (25) that rate-equation models neglecting intracellular dynamics overestimate population growth. Intuitively, the reason is that observed response-time distributions fall in between exponential response-time distributions and deterministic doubling times, corresponding to a doubling time scaling with base *e* vs. base 2. Furthermore, we analysed the robustness of regulatory mechanisms for cell-growth control. Previous models of lymphocyte dynamics have often accounted for such regulation by considering time dependencies of the cell-division rate *β* = *β*(*t*) (27,54), and we sought to derive a plausible mechanistic formulation of that time dependence. We found that a combination of an IL-2 dependent quorum sensing type mechanism, together with a previously described genetic timer mechanism (37), had valuable properties regarding tunability and robustness also to repetitive stimulation such as in chronic inflammation scenarios.

Chronic inflammation can occur in a spectrum of human diseases including viral infections such as hepatitis, autoimmune conditions such as rheumatoid arthritis and the tumour microenvironment in various cancer entities (9,11). How and why chronic inflammation occurs is incompletely understood, but a common rational is that chronification and thus stabilization of a low-grade inflammation may be a suitable means to avoid overshooting immune responses and autoimmune organ damage in the case of severe infections (42). Combining response-time modeling with an annotation strategy based on high-resolution in vitro data and specific ex vivo data allowed us to derive a model that reflects the expected stabilization properties of chronic inflammation and is constrained to a narrow range of dynamic behaviour. Notably, the relapse-effects predicted by our model for a chronic inflammation scenario are commonly found in several autoimmune disorders (9).

While several factors have been identified that contribute to the emergence and stabilization of a chronic infection, our model simulations show that two data-derived constraints, namely negative feedback via IL-10 together with antigen-driven proliferation of precursor cells (34,46,47), are sufficient to capture the experimentally observed chronic Th cell response. Further, our model simulations show that the discovery of intermediate proliferative precursor cells that persist during chronic viral infection (34,46) is well in line with previously described infection kinetics (51). Similar progenitor-like phenotypes have been found for exhausted CD8+ T cells (34), thus indicating a general motif of immunological decision-making.

Optimal treatment timing is a prevailing question in clinical therapy development. For instance, early treatment of rheumatoid arthritis is a key predictor of disease remission, and similar “hit hard and early” strategies have been suggested for treatment of multiple sclerosis and systemic lupus erythematosus (55). However, in many cases longitudinal data to systemically identify time-dependent therapeutic effect sizes are sparse, which confounds precise estimates of optimal intervention times. Here, performing systematic in silico analyses of the fully annotated model revealed a tongue-shaped landscape of optimal intervention times. In fact, our simulations rather point towards a transient window of opportunity for optimal efficacy of an intervention. That window of opportunity sensitively depends on the specific situation, and we anticipate that identification of such windows of opportunities will be an important component of effective personalized therapy strategies in the future.

Overall, our simulations highlight a network view on immune-cell decision making. That is in line with recently performed detailed comparisons of immune responses against infections with genetically similar virus strains, namely influenza vs. mild or severe *Covid-19* (56,57), which revealed subtle but widespread differences in various cytokine species that go along with large differences in the strength of the immune response. The mathematical framework and the data-driven model of Th cell dynamics presented here may lay the foundation for further research towards quantitative understanding of immune-cell decision circuits.

## Methods

### Integrated response-time modelling framework

Consider a cell type in which each cell can take a set of distinct cellular states *i* = 1,…,*n* and let *S*_1_(*t*),…*S_n_*(*t*) denote the number cells in each state at time *t*. Further, suppose that the system evolves in time by cell-state transitions *i* → *j* that occur at times drawn from a response time probability distribution *Ψ_ij_*(*t-τ*), where *τ* is the time of the last state-change in the system. Of note, *Ψ_ij_*(*t - τ*) is often called a first-passage time distribution in physics (30), or a phase-type distribution in the mathematical literature (58). In the following, we restrict ourselves to cases where *Ψ_ij_*(*t - τ*) takes the form of an Erlang distribution

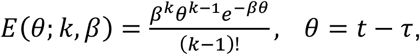

which is the special case of a gamma distribution with integer-valued shape parameter *k*. Hence, we set *Ψ_ij_*(*t - τ*) = *E*(*t - τ;k_ij_,β_ij_*), and in that case, the average dynamics of the system are equivalent to a set of ordinary differential equations that can be derived using the linear-chain trick (Figure S1A-C) (24,25,59). Following (59), we reduce the system by assuming that the time to remain in state *i* is independent of the decision process *i* → *j*, that is, the probability *p_ij_* of the transition *i* → *j* is an additional parameter that is independent of the response-time distribution. Consequently, *Ψ_ij_*(*t - τ*) is reduced to an Erlang distribution *E*(*t - τ;k_i_,β_i_*) that only depends on the cell-state of origin. We further account for cell-cell communication (24) in the form of cell-state transitions that are modulated by the concentrations of cytokine species *c_m_,m* = 1 …*M*, which are produced and taken up with cellstate specific rates. We describe cell proliferation by assuming that with probability *p_ii_*, cells will give rise to *d_i_* cells in state *i* rather than transitioning to a new state *j*, where *d_i_* is the division number (25,60).

Combining those processes gives rise to a closed mathematical framework that allows for incorporating branched decision-making and cell proliferation, in addition to cell-cell-communication dynamics:

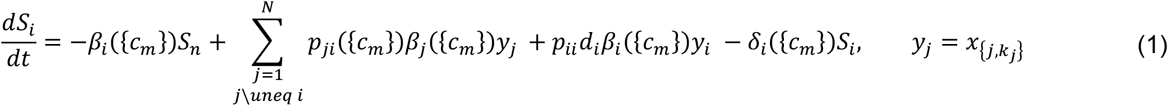

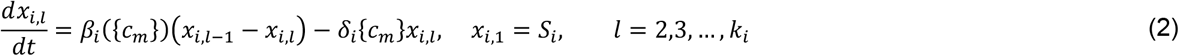

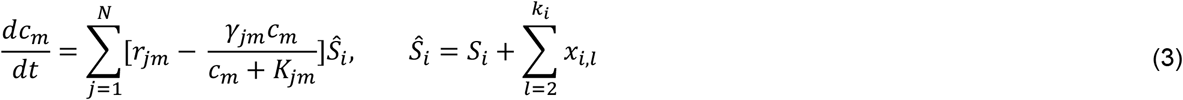

Here, the last term in Equation 1 accounts for cell death at rate *δ_i_*({*c_m_*}), in addition to the processes described above. In our case, the response-time distributions for differentiation and proliferation obtained from data are similar enough to be well described by only one Erlang distribution (Figure S2D), so that both processes are treated by a single multi-step process (Equation 2). Otherwise, the framework can easily be generalized to separate distributions by taking into account an additional, competing chain 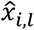 for proliferation, supplemented by exchange processes between the differentiation and proliferation chains (59). Regarding cytokine dynamics (Equation 3), we consider that cells may have a limited uptake capacity, represented by a set of half-saturation constants *K_jm_* (38). The cytokine concentrations {*c_m_*} can effect both reaction times and reaction probabilities in every cell-state by means of a state-function 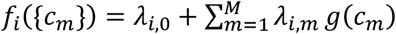, where for *g*(*c_m_*) we either chose Hill-type kinetics taking the form 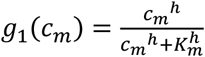 or a fold-change variant 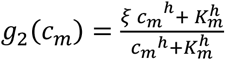 where *ξ* represents the feedback fold-change. Further, we take 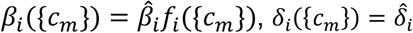 and 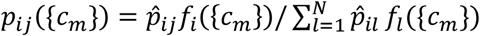, the latter ensuring normalization of the branching probabilities such that 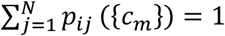.

Network motif models, proliferation models and the chronic inflammation model were all derived based on Equations 1–3, the full equations for the respective models are displayed in the Supplementary Text.

### Kinetic transcriptome analysis

To analyze kinetic T cell genes, we used published microarray and RNA-seq data (45,60–63) (Table 2). Expression values were normalized using standard methods and kinetic genes were identified with a regression-based as described previously (62). To further normalize the data for response-time modeling, the expression values were min-max normalized for each gene across time points. We defined two categories of kinetic patterns based on the shape parameter *α* of the gamma distribution *γ*(*α,β*): Model A (*α* > 1, delayed onset kinetics) and Model B (*α* = 1, exponential onset kinetics). The cumulative gamma distribution *F*(*x,α,β*) of the defined models was fitted for each gene in the data sets to identify distribution parameters *α* and *β* (Model A, B) or only *β* (Model B) by non-linear least squares fitting. For model selection, we used an F-test for nested models with correction for multiple testing. Genes with poor fit quality (RMSE > 0.3) were assigned to the “other” category. In this last category, we additionally assessed bimodality by fitting means *μ*_1_, *μ*_2_ and standard deviations *σ*_1_, *σ*_2_ of the CDF of a two-component Gaussian mixture model *F*(*x*) = *w*_1_*Φ*_1_(*x*, *μ*_1_, *σ*_1_) + *w*_2_*Φ*_2_(*x*,*μ*_2_,*σ*_2_) and assigned a gene as bimodal if RMSE < 0.3. Equal standard deviations and weights were assumed (*w*_1_ = *w*_2_,*σ*_1_ = *σ*_2_).

**Table 2:** Studies used for data analysis.

| Study | Cell Types | ID | Source |
| --- | --- | --- | --- |
| Crawford et al. 2014 | CD4+, CD8+ T cells | GSE30431 | (45) |
| Ilott et al. 2016 | Bulk Lymph Node | E-MTAB-3590 | (61) |
| Proserpio et al. 2016 | Th2 cells | E-MTAB-3543 | (60) |
| Burt et al. 2022 | Th1,Th2, Th1/2, Th0 cells | GSE200250 | (62) |
| Nir et al. 2013 | Th0, Th17 cells | GSE163345 | (63) |

### Gene module analysis and data integration

Gene lists for Th1 and Th2 differentiation, as well as cytokines, transcription factors and cytokine receptor gene lists were taken from Stubbington et al. (32). Other gene modules were derived from the GO:BP category from the Msigdbr data base. Data for T-bet and Gata-3 protein kinetics were taken from Peine et al. (64) (cf. Figure 2D). To assign a gene to a specific category (gamma, expo, other, bimodal) across several data sets, we counted the number of fit assignments, and the gene was assigned to the category with the maximum number of assignments. Only genes were considered that were found in at least 5 data sets. Significance testing for enrichment analysis was performed by applying a hypergeometric test with subsequent correction for multiple testing using the clusterProfiler package in R.

### Numerical simulations and Statistics

ODE models were solved numerically using the LSODA algorithm (Adams/BDF solvers) implemented in the SciPy library in python. For simulations with repeated stimulation, the ODE system was integrated in a piece-wise fashion with initial values being updated at each event. For curve-fitting of the chronic inflammation model, we used least-squares optimization (Levenberg–Marquardt algorithm) as implemented in the Python lmfit library. Statistical tests were performed with correction for multiple testing using the Benjamini-Hochberg method at false-discovery-rate below 10%.

## Acknowledgements

We thank Max Löhning, Caroline Peine, Valerie Plajer and members of the Thurley group for fruitful discussions, and Helene Glöckner for assistance in implementation of model simulations. This work was supported by the Deutsche Forschungsgemeinschaft (TH 1861/4-1), Germany’s Excellence Strategy (EXC2151 and EXC2047, to K.T.), the Leibniz Association (Best-minds junior research group, to K.T.) and the Joachim Herz Stiftung (to P.B.).

## References

1. Cohen M, Giladi A, Barboy O, Hamon P, Li B, Zada M, Gurevich-Shapiro A, Beccaria CG, David E, Maier BB, et al. The interaction of CD4+ helper T cells with dendritic cells shapes the tumor microenvironment and immune checkpoint blockade response. Nat cancer (2022) 3:303–317.

2. Rao DA, Gurish MF, Marshall JL, Slowikowski K, Fonseka CY, Liu Y, Donlin LT, Henderson LA, Wei K, Mizoguchi F, et al. Pathologically expanded peripheral T helper cell subset drives B cells in rheumatoid arthritis. Nature (2017) 542:110–114.

3. Schrier SB, Hill AS, Plana D, Lauffenburger DA. Synergistic Communication between CD4+ T Cells and Monocytes Impacts the Cytokine Environment. Sci Rep (2016) 6:34942.

4. Chua RL, Lukassen S, Trump S, Hennig BP, Wendisch D, Pott F, Debnath O, Thürmann L, Kurth F, Völker MT, et al. COVID-19 severity correlates with airway epithelium– immune cell interactions identified by single-cell analysis. Nat Biotechnol (2020) 38:970–979.

5. Arnold KB, Szeto GL, Alter G, Irvine DJ, Lauffenburger DA. CD4+ T cell-dependent and CD4+ T cell-independent cytokine-chemokine network changes in the immune responses of HIV-infected individuals. Sci Signal (2015) 8:ra104.

6. Barry KC, Hsu J, Broz ML, Cueto FJ, Binnewies M, Combes AJ, Nelson AE, Loo K, Kumar R, Rosenblum MD, et al. A Natural Killer/Dendritic Cell Axis Defines Responsive Tumor Microenvironments in Melanoma. Nat Med (2018) 24:1178.

7. Burmester GR, Feist E, Dörner T. Emerging cell and cytokine targets in rheumatoid arthritis. Nat Rev Rheumatol (2014) 10:77–88.

8. Altan-Bonnet G, Mukherjee R. Cytokine-mediated communication: a quantitative appraisal of immune complexity. Nat Rev Immunol (2019) 19:205–217.

9. Maschmeyer P, Chang HD, Cheng Q, Mashreghi MF, Hiepe F, Alexander T, Radbruch A. Immunological memory in rheumatic inflammation - a roadblock to tolerance induction. Nat Rev Rheumatol (2021) 17:291–305.

10. Schwartz DM, Bonelli M, Gadina M, O’Shea JJ. Type I/II cytokines, JAKs, and new strategies for treating autoimmune diseases. Nat Rev Rheumatol (2015) 12:25–36.

11. Briukhovetska D, Dörr J, Endres S, Libby P, Dinarello CA, Kobold S. Interleukins in cancer: from biology to therapy. Nat Rev Cancer (2021) 21:481–499.

12. Morel PA, Lee REC, Faeder JR. Demystifying the cytokine network: Mathematical models point the way. Cytokine (2016) 98:115–123.

13. Heinrich R, Neel BG, Rapoport TA. Mathematical models of protein kinase signal transduction. Mol Cell (2002) 9:957–70.

14. Alon U. Network motifs: theory and experimental approaches. Nat Rev Genet 2007 86 (2007) 8:450–461.

15. El-Samad H. Biological feedback control—Respect the loops. Cell Syst (2021) 12:477–487.

16. Pritchard JR, Bruno PM, Gilberta LA, Caprona KL, Lauffenburger DA, Hemann MT. Defining principles of combination drug mechanisms of action. Proc Natl Acad Sci U S A (2013) 110:E170–E179.

17. Klinger B, Sieber A, Fritsche-Guenther R, Witzel F, Berry L, Schumacher D, Yan Y, Durek P, Merchant M, Sch??fer R, et al. Network quantification of EGFR signaling unveils potential for targeted combination therapy. Mol Syst Biol (2013) 9: 673.

18. Saez-Rodriguez J, Blüthgen N. Personalized signaling models for personalized treatments. Mol Syst Biol (2020) 16:e9042.

19. Behar M, Barken D, Werner SL, Hoffmann A. The dynamics of signaling as a pharmacological target. Cell (2013) 155: 448–61.

20. Yofe I, Dahan R, Amit I. Single-cell genomic approaches for developing the next generation of immunotherapies. Nat Med (2020) 26:171–177.

21. Haque A, Engel J, Teichmann SA, Lönnberg T. A practical guide to single-cell RNA-sequencing for biomedical research and clinical applications. Genome Med(2017) 9:1–12.

22. Luecken MD, Theis FJ. Current best practices in single-cell RNA-seq analysis: a tutorial. Mol Syst Biol (2019) 15:e8746

23. Nienałtowski K, Rigby RE, Walczak J, Zakrzewska KE, Głów E, Rehwinkel J, Komorowski M. Fractional response analysis reveals logarithmic cytokine responses in cellular populations. Nat Commun 2021 121 (2021) 12:1–10.

24. Thurley K, Wu LF, Altschuler SJ. Modeling Cell-to-Cell Communication Networks Using Response-Time Distributions. Cell Syst (2018) 6:355–367.e5.

25. Yates CA, Ford MJ, Mort RL. A Multi-stage Representation of Cell Proliferation as a Markov Process. Bull (2017) 79:2905–2928.

26. Castro M, López-García M, Lythe G, Molina-París C. First passage events in biological systems with non-exponential inter-event times. Sci Rep (2018) 8:

27. De Boer RJ, Perelson AS. Quantifying T lymphocyte turnover. J Theor Biol (2013) 327:45–87.

28. Kim DW, Hong H, Kim JK. Systematic inference identifies a major source of heterogeneity in cell signaling dynamics: The rate-limiting step number. Sci Adv (2022) 8:4598.

29. Choi B, Cheng YY, Cinar S, Ott W, Bennett MR, Josić K, Kim JK. Bayesian inference of distributed time delay in transcriptional and translational regulation. Bioinformatics (2020) 36:586–593.

30. Kampen VNG. Stochastic Processes in Physics and Chemistry. 3rd ed. North-Holland Personal Library (2007).

31. Zhu J, Yamane H, Paul WE. Differentiation of Effector CD4 T Cell Populations. Annu Rev Immunol (2010) 28:445–489.

32. Stubbington MJT, Mahata B, Svensson V, Deonarine A, Nissen JK, Betz AG, Teichmann SA. An atlas of mouse CD4+ T cell transcriptomes. Biol Direct (2015) 10:

33. Hart Y, Antebi YE, Mayo AE, Friedman N, Alon U. Design principles of cell circuits with paradoxical components. Proc Natl Acad Sci U S A (2012) 109:8346–51.

34. Chu HH, Chan SW, Gosling JP, Blanchard N, Tsitsiklis A, Lythe G, Shastri N, Molina-París C, Robey EA. Continuous Effector CD8+T Cell Production in a Controlled Persistent Infection Is Sustained by a Proliferative Intermediate Population. Immunity (2016) 45:159–171.

35. Gett A V, Hodgkin PD. A cellular calculus for signal integration by T cells. Nat Immunol (2000) 1:239–244.

36. Hammer Q, Rückert T, Borst EM, Dunst J, Haubner A, Durek P, Heinrich F, Gasparoni G, Babic M, Tomic A, et al. Peptide-specific recognition of human cytomegalovirus strains controls adaptive natural killer cells. Nat Immunol (2018) 19:1–11.

37. Heinzel S, Binh Giang T, Kan A, Marchingo JM, Lye BK, Corcoran LM, Hodgkin PD. A Myc-dependent division timer complements a cell-death timer to regulate T cell and B cell responses. Nat Immunol (2017) 18:96–103.

38. Zhou X, Franklin RA, Adler M, Jacox JB, Bailis W, Shyer JA, Flavell RA, Mayo A, Alon U, Medzhitov R. Circuit Design Features of a Stable Two-Cell System. Cell (2018) 172:744–757.e17.

39. Almeidal ARM, Amado IF, Reynolds J, Berges J, Lythe G, Molina-París C, Freitas AA. Quorum-sensing in CD4+ T cell homeostasis: A hypothesis and a model. Front Immunol (2012) 3:125.

40. Huang J, Brameshuber M, Zeng X, Xie J, Li Q jing, Chien Y hsiu, Valitutti S, Davis MM. A Single peptide-major histocompatibility complex ligand triggers digital cytokine secretion in CD4+ T Cells. Immunity (2013) 39:846–857.

41. Vella AV, Herati SR, Wherry EJ. The Tfh perspective. Trends Mol Med (2017) 23:1072–1087.

42. Wherry EJ, Kurachi M. Molecular and cellular insights into T cell exhaustion. Nat Rev Immunol (2015) 15:486–499.

43. Brooks DG, Teyton L, Oldstone MBA, McGavern DB. Intrinsic Functional Dysregulation of CD4 T Cells Occurs Rapidly following Persistent Viral Infection. J Virol (2005) 79:10514–10527.

44. Matloubian M, Concepcion RJ, Ahmed R. CD4+ T cells are required to sustain CD8+ cytotoxic T-cell responses during chronic viral infection. J Virol (1994) 68:8056–8063.

45. Crawford A, Angelosanto JM, Kao C, Doering TA, Odorizzi PM, Barnett BE, Wherry EJ. Molecular and Transcriptional Basis of CD4+ T Cell Dysfunction during Chronic Infection. Immunity (2014) 40:289–302.

46. Xia Y, Sandor K, Pai JA, Daniel B, Raju S, Wu R, Hsiung S, Qi Y, Yangdon T, Okamoto M, et al. BCL6-dependent TCF-1+ progenitor cells maintain effector and helper CD4+ T cell responses to persistent antigen. Immunity (2022) 55:1200–1215.e6.

47. Snell LM, Osokine I, Yamada DH, De la Fuente J, Elsaesser HJ, Elsaesser HJ, Brooks DG. Overcoming CD4 Th1 Cell Fate Restrictions to Sustain Antiviral CD8 T Cells and Control Persistent Virus Infection. Cell Rep (2016) 16:3286–3296.

48. Polonsky M, Rimer J, Kern-Perets A, Zaretsky I, Miller S, Bornstein C, David E, Kopelman NM, Stelzer G, Porat Z, et al. Induction of CD4 T cell memory by local cellular collectivity. Science (2018) 360:

49. Zaretsky I, Polonsky M, Shifrut E, Reich-Zeliger S, Antebi Y, Aidelberg G, Waysbort N, Friedman N. Monitoring the dynamics of primary T cell activation and differentiation using long term live cell imaging in microwell arrays. Lab Chip (2012) 12:5007.

50. Löhning M, Hegazy AN, Pinschewer DD, Busse D, Lang KS, Höfer T, Radbruch A, Zinkernagel RM, Hengartner H. Long-lived virus-reactive memory T cells generated from purified cytokine-secreting T helper type 1 and type 2 effectors. J Exp Med (2008) 205:53–61.

51. Fahey LM, Wilson EB, Elsaesser H, Fistonich CD, McGavern DB, Brooks DG. Viral persistence redirects CD4 T cell differentiation toward T follicular helper cells. J Exp Med (2011) 208:987–999.

52. Suter DM, Molina N, Gatfield D, Schneider K, Schibler U, Naef F. Mammalian genes are transcribed with widely different bursting kinetics. Science (2011) 332:472–4.

53. Stumpf PS, Smith RCG, Lenz M, Schuppert A, Müller FJ, Babtie A, Chan TE, Stumpf MPH, Please CP, Howison SD, et al. Stem Cell Differentiation as a Non-Markov Stochastic Process. Cell Syst (2017) 5:268–282.e7.

54. Callard R, Hodgkin P. Modeling T-and B-cell growth and differentiation. (2007). Immunol Rev 216:119–29.

55. Van Nies JAB, Tsonaka R, Gaujoux-Viala C, Fautrel B, Van Der Helm-Van Mil AHM. Evaluating relationships between symptom duration and persistence of rheumatoid arthritis: Does a window of opportunity exist? Results on the Leiden Early Arthritis Clinic and ESPOIR cohorts. Ann Rheum Dis (2015) 74:806–812.

56. Youngs J, Provine NM, Lim N, Sharpe HR, Amini A, Chen YL, Luo J, Edmans MD, Zacharopoulou P, Chen W, et al. Identification of immune correlates of fatal outcomes in critically ill COVID-19 patients. PLOS Pathog (2021) 17:e1009804.

57. Karagiannis F, Peukert K, Surace L, Michla M, Nikolka F, Fox M, Weiss P, Feuerborn C, Maier P, Schulz S, et al. Impaired ketogenesis ties metabolism to T cell dysfunction in COVID-19. Nature (2022) 609:801–807.

58. Bladt M. A Review on Phase-type Distributions and their Use in Risk Theory. Astin Bull (2005) 35:145–161.

59. Hurtado PJ, Kirosingh AS. Generalizations of the ‘Linear Chain Trick’: incorporating more flexible dwell time distributions into mean field ODE models. J Math Biol (2019) 79:1831–1883.

60. Proserpio V, Piccolo A, Haim-Vilmovsky L, Kar G, Lönnberg T, Svensson V, Pramanik J, Natarajan KN, Zhai W, Zhang X, et al. Single-cell analysis of CD4+ T-cell differentiation reveals three major cell states and progressive acceleration of proliferation. Genome Biol (2016) 17:103.

61. Ilott NE, Bollrath J, Danne C, Schiering C, Shale M, Adelmann K, Krausgruber T, Heger A, Sims D, Powrie F. Defining the microbial transcriptional response to colitis through integrated host and microbiome profiling. (2016) 10:2389–2404.

62. Burt P, Peine M, Peine C, Borek Z, Serve S, Floßdorf M, Hegazy AN, Höfer T, Löhning M, Thurley K. Dissecting the dynamic transcriptional landscape of early T helper cell differentiation into Th1, Th2, and Th1/2 hybrid cells. Front Immunol (2022) 13:1–11.

63. Yosef N, Shalek AK, Gaublomme JT, Jin H, Lee Y, Awasthi A, Wu C, Karwacz K, Xiao S, Jorgolli M, et al. Dynamic regulatory network controlling TH17 cell differentiation. Nature (2013) 496:461–468.

64. Peine M, Rausch S, Helmstetter C, Fröhlich A, Hegazy AN, Kühl AA, Grevelding CG, Höfer T, Hartmann S, Löhning M. Stable T-bet+GATA-3+ Th1/Th2 Hybrid Cells Arise In Vivo, Can Develop Directly from Naive Precursors, and Limit Immunopathologic Inflammation. PLoS Biol (2013) 11:e1001633.

