## Supplemental Material for "Distribution-modeling quantifies collective Th cell decision circuits in chronic inflammation"

### Supplementary Material

#### 1 Supplementary Text

In the following, the mathematical models introduced in the main text are specified. All state-transitions shown below were translated into differential rate-equations using our proposed framework (cf. Equations 1-2, main text). All cytokine dynamics were modeled according to Equation 3, main text.

##### Minimal branching models

The core motif (Figure 1A) consists of a model where an activated cell ThN could differentiate into two cell types ThX and ThY. Let the set $S = \{S_x, S_y, S_n\}$ denote cell types $\{\text{ThN}, \text{ThX}, \text{ThY}\}$ respectively, then we get:

$$\begin{aligned}
S_n &\xrightarrow{E(k_{S_n},\beta_{S_n})} p_{S_x} S_x \\
S_x &\xrightarrow{E(k_{S_n},\beta_{S_n})} p_{S_y} S_y \\
S_x &\xrightarrow{E(k_{S_x},\beta_{S_x})} 2S_x \\
S_y &\xrightarrow{E(k_{S_y},\beta_{S_y})} 2S_y \\
S_x &\xrightarrow{\delta_{S_x}} \emptyset \\
S_y &\xrightarrow{\delta_{S_y}} \emptyset,
\end{aligned} \tag{1}$$

where $E(k, \beta)$ represents the respective Erlang-distribution for each state transition, $p_x$ and $p_y$ represent the branching probabilities and $\delta_i$ the death rates. To account for feedback (cf. Figure 1A, model ii), we modeled cytokines $c_1$ and $c_2$ assuming feedback on the respective branching probability $p_x$ and $p_y$.

##### Cytokine blockade model

For the cytokine blockade model (Figure 1A, model iii), we did not model cytokine dynamics explicitly and instead modulated the branching probability directly within a given time-window using the step-function approximation:

$$p_{S_x}(t) = 0.5 \cdot \left[p_w + \hat{p}_{S_x} + (p_w - \hat{p}_{S_x}) \tanh(s(t - t_1)) \tanh(s(t_2 - t))\right] \tag{2}$$

where $t_1$ and $t_2$ denote the first and last times for the blockade (cf. Figure 1, main text), $p_w$ denotes the fold-change in probability within the decision window and $s$ denotes the steepness of the window with higher values leading to a more step-like window. Effect sizes for early, intermediate and late intervention times were derived by computing the maximum ThX-to-ThY log2 fold-change for individual simulations compared to a control scenario without cytokine blockade with parameter $\hat{p}_{S_x}$ being drawn from a uniform distribution $U(0, 1)$.

#### Parameter values

Since we were here interested in generic network properties, we did not use any data-estimates for parameter annotation and instead tried to keep the network topology and parameters as simple as possible: $k_{S_n} = k_{S_y} = k_{S_x} = 10$, $\beta_{S_n} = \beta_{S_x} = \beta_{S_y} = 10.0$ d$^{-1}$ (RTM model), $k_{S_n} = k_{S_y} = k_{S_x} = 1$, $\beta_{S_n} = \beta_{S_x} = \beta_{S_y} = 1.0$ d$^{-1}$ (SSM model). Branching probabilities: $\hat{p}_{S_X} = \hat{p}_{S_y} = 0.5$. Cell death: $\delta_{S_x} = \delta_{S_y} = 2$ d$^{-1}$. Cytokine secretion: $r_{S_x,1} = r_{S_y,2} = 1$ molecules $s^{-1}$ cell$^{-1}$. Cytokine uptake: $\gamma_{S_x,1} = \gamma_{S_y,2} = 1 s^{-1}$ cell$^{-1}$. Feedback: $\xi_{S_x} = \xi_{S_y} = 10$, $p_w = 10$, $s = 100$.

#### Proliferation models

The models in Figure 3A (main text) were based on a 3-state-model where activated naive cells $S_n$ transition into precursor cells $S_p$ and finally into proliferating effector cells $S_e$, resulting in the following system:

$$\begin{aligned} S_n &\xrightarrow{E(k_{S_n},\beta_{S_n})} S_p \\ S_p &\xrightarrow{E(k_{S_p},\beta_{S_p})} 2S_e \\ S_e &\xrightarrow{E(k_{S_e},\beta_{S_e})} 2S_e \\ S_e &\xrightarrow{\delta_{S_e}} \emptyset, \end{aligned} \tag{3}$$

#### Carrying capacity model

For the carrying capacity model, we assumed: $\beta(S_e) = \begin{cases} \hat{\beta}_e, & \text{if } \hat{S}_e < C \\ 0, & \text{else,} \end{cases}$ where $C$ denotes the carrying capacity.

#### Timer Model

For the Timer model, we assumed exponential decay of Myc:

$$\frac{d\text{Myc}}{dt} = -\delta_{\text{Myc}}[\text{Myc}], \tag{4}$$

where $\delta_{[\text{Myc}]}$ denotes Myc degradation.

#### IL2 model

For the IL-2 and Timer models, we assumed regulation of the form $\beta_e(x) = \hat{\beta}_e \cdot f(x)$ with $f(x)$ representing a Michaelis-Menten-type function $f(x) = x^3/(x^3 + K^3)$ where $x$ would represent either IL-2 or the transcription factor Myc (Timer Model) and $K$ the corresponding half-saturation constant. Generally, we assumed that activated Naive and Precursor cells were the major source of IL-2 and thus only considered effective secretion of those cell types.

#### Mixed Timer and IL2 model

In case of the mixed Timer and IL-2 model we assumed multiplicative signal integration of the form $\beta_e(x) = \hat{\beta}_e \cdot f(\mathrm{IL2})f(\mathrm{Myc})$.

#### Restimulation

For restimulation experiments, we also considered secretion of IL-2 by effector cells. In this case, it was assumed that for a set of restimulations $\{\tau_1, ..., \tau_n\}$, the secretion rate declined exponentially after each restimulation event:

$$r_{\mathrm{IL2}}(t,\tau) = \begin{cases} 0, & t < \tau_1 \\ r_{\mathrm{IL2}} \cdot e^{\delta_{\mathrm{IL2}}(t-\tau_i)}, & \tau_i \leq t < \tau_{i+1}, \quad i \in \{1,2,...,\tau_{n-1}\}, \end{cases} \tag{5}$$

where $t$ denotes time, $\tau$ is the time of restimulation and $\delta_{\mathrm{IL2}}$ is the decay rate for IL-2 secretion after restimulation. Restimulation events were assumed to be exponentially distributed with rate parameter $\lambda$. The inverse CDF of the exponential distribution was used to compute restimulation times with random sampling from a uniform distribution $U(0,1)$:

$$F_X^{-1}(U) = -\frac{ln(1-U)}{\lambda}. \tag{6}$$

#### Parameter values

$C = 1000$ cells, $\delta_{\mathrm{IL2}} = 2\ \mathrm{d}^{-1}$, $\lambda = 0.1\ \mathrm{d}^{-1}$. IL-2 secretion: $r_{\mathrm{IL2}} = 100$ molecules cell$^{-1}$ $s^{-1}$. State transitions: $k_{S_n} = 22$, $\beta_{S_n} = 15.0$, $k_{S_p} = k_{S_e} = 7$, $\beta_{S_p} = \beta_{S_e} = 15.2$. The remaining model specific parameters ($\delta_{\mathrm{Myc}} = 0.37\ \mathrm{d}^{-1}$, $\gamma_{\mathrm{IL2}} = 4.5$ molecules cell$^{-1}$ $s^{-1}$) were adjusted to achieve the same peak height as the carrying capacity model (cf. Figure 3B, main text).

#### Modeling Th cell dynamics in acute and chronic infection

To derive a model of in vivo Th cell dynamics observed for chronic and acute viral infection, we extended the data-annotated proliferation model by adding effector subtypes, chronic and memory cells. Briefly, activated, naive cells ($A_1$) transition into precursor cells ($A_2$). We assumed that with a given probability the precursor cells will remain in the precursor state upon division instead of entering further differentiation programs. The precursor cells can develop into terminally differentiated effector cells through a course of intermediary transitions. Terminally differentiated cells can adopt memory or chronic phenotypes. Let

$X \in \{\text{Th1}, \text{Tfh}\}$ denote the two effector cell types Th1 and Tfh. We then get:

$$\begin{aligned}
A_1 &\xrightarrow{E(k_{a_1},\beta_{a_1})} 2A_2 \\
A_2 &\xrightarrow{E(k_{a_2},\beta_{a_2})} 2p_{A_2}A_2 \\
A_2 &\xrightarrow{E(k_{a_2},\beta_{a_2})} 2p_X X^1_{\text{int}}, \quad X \in \{\text{Th1}, \text{Tfh}\} \\
X^i_{\text{int}} &\xrightarrow{E(k_e,\beta_e)} 2X^{i+1}_{\text{int}}, \quad i \in \{1,2,3\}, X^4_{\text{int}} = X_t \\
X_t &\xrightarrow{\beta_m} X_m \\
X_t &\xrightarrow{\beta_c} X_c
\end{aligned} \tag{7}$$

where the subscripts $\{\text{int}, t, m, c\}$ indicate intermediate effector cells, terminally differentiated cells, memory cells and chronic cells respectively. We additionally modeled cell death based on a rate process assuming that cells $A_2, X^i_{\text{int}}, X_t, X_t$ died with decay rate $\delta_e$. Memory cell turnover was modeled using decay rate $\delta_m$ and for naive Th cells we assumed that cell death was negligible compared to differentiation. Further, we assumed that divisions of intermediate cell types followed the same distribution as the proliferating precursor cell population so that $\beta_e = \beta_{a_2}$ and $k_e = k_{a_2}$. Antigen dynamics were modeled assuming exponential decay of the antigen term:

$$\frac{d[\text{ag}]}{dt} = -\delta_{\text{ag}} \cdot [\text{ag}] \tag{8}$$

For the branching probabilities $p_{\text{Th1}}$, and $p_{\text{Tfh}}$, we assumed modulation through antigen and IL-10 in the form

$$p_i(\text{ag}, \text{IL-10}) = \lambda_{i,0} + \lambda_{i,\text{ag}} \cdot f(\text{ag}) + \lambda_{i,\text{IL10}} \cdot f(\text{IL-10}), \quad i \in \{\text{Th1}, \text{Tfh}\}, \tag{9}$$

and for proliferation, we built on the previously established model but added antigen and IL-10 effects:

$$\beta_e(\text{ag}, \text{IL-10}, \text{IL-2}, \text{Myc}) = \hat{\beta}_e \left[\lambda_{e,\text{ag}} f(\text{ag}) + \lambda_{e,\hat{\text{ag}}} f(\text{IL-2}) f(\text{Myc}) g(\text{IL-10})\right] \tag{10}$$

In Equations 9 and 10, $f$ denotes a Hill-type function of the form $f(c) = c^3/(c^3 + K^3)$ and we used $g(c) = (\xi c^3 + K^3)/(c^3 + K^3)$ to model negative IL-10 feedback with $\xi < 1$ representing the feedback fold-change. To ensure that the maximum proliferation rate is $\hat{\beta}_e$, we further add the normalization constraint $\lambda_{e,\text{ag}} + \lambda_{e,\hat{\text{ag}}} = 1$. For the branching probabilities, we impose that $\sum_i \lambda_{i,0} = 1$, with $p_{A_2} = \lambda_{A_2,0}$. For a summary of parameter descriptions and values, refer to Table 2, and Table S3.

#### 2 Supplementary Tables

##### Table S1: Fit-statistics derived from kinetic transcriptome analysis.

Each row represents the fit statistics root-sum-squares (rss) and root-mean-square error (rmse), along with standard error estimates for parameters, for an individual gene for a fit to an exponential or gamma distribution. The "column model" indicates the best fit model. Tabs represent individual data sets and conditions (separate .xlsx file).

##### Table S2: Overrepresentation analysis of gene sets across different fit categories.

For the pathway analysis, genes in the gamma- and exponential categories from the data analysis were used (cf. Figure 2). Shown are original and FDR-corrected p-values for each pathway. Gene sets were derived from the REACTOME data base. Tabs represent enrichment analyses for individual data sets and conditions (cf. Table 2) (separate .xlsx file).

#### Table S3: Additional parameter values

| Symbol | Parameter | Value | Unit | Source |
|---|---|---|---|---|
| $r_{IL2}$ | Rate of IL-2 secretion | 10 | Molecules $cell^{-1}s^{-1}$ | [47] |
| $\gamma_{IL2}$ | Rate of IL-2 uptake | 1 | Molecules $cell^{-1}s^{-1}$ | [47] |
| $K_{IL10}$ | Half Saturation for IL-10 | 18.6 | pM | [8] |
| $K_{IL2}$ | Half Saturation for IL-2 | 15.5 | pM | [8] |
| $r_{IL10,e}$ | Rate of IL-10 secretion by Th1 effector cells | 10 | Molecules $cell^{-1}s^{-1}$ | - |
| $r_{IL10,c}$ | Rate of IL-10 secretion by chronic Th1 cells | 100 | Molecules $cell^{-1}s^{-1}$ | - |
| $\lambda_{Th1,IL10}$ | IL-10 effect branching probability Th1 | 10 | - | - |
| $\lambda_{Tfh,ag}$ | Antigen effect branching probability Th1 | 0.5 | - | - |
| $\delta_{Myc}$ | Degradation rate Myc | 0.37 | 1/d | - |
| $\delta_{ag}$ | Degradation rate antigen | 0.1 | 1/d | - |
| $\delta_{IL10}$ | Global IL-10 consumption | $5\ 10^4$ | Molecules $s^{-1}$ | - |
| $K_{Myc}$ | Half Saturation Timer | 0.1 | a.u. | - |
| $K_{ag}$ | Half Saturation for antigen affecting proliferation | 0.5 | a.u. | - |
| $n_{Tregs}$ | Number of regulatory T cells | 1000 | - | - |

#### 3 Supplementary Figure Legends

##### Figure S1: Supplementary data to Figure 1.

**(A)** Extended model scheme (cf. Figure 1A). Cell X may differentiate into cell Y or Z, or remain in state X and proliferate (cf. Methods). If the time to remain in state X is well described by an Erlang-distribution with shape parameter k, then the process can be described as a linear chain with k steps (cf. Equations 1-2, main text). **(B)** Shape of the Erlang-distribution for different values of the parameter k. **(C)** Stochastic simulation of the model in (A). Shown are mean (solid lines) and s.d. for 50 simulations with 50 cells per run. **(D-G)** Response-time modeling with competing state-transitions. **(D)** model scheme for a binary cell-fate decision based on competing response-time distributions.**(E)** Model simulations , for scenarios with equal means $\mu$ but different chain lengths (left) or with unequal means $\mu$ and different chain lengths (right). **(F)** Feedback effects during competition. Shown is the Y-to-Z ratio as a function of the feedback fold-change in a scenario with equal means but different chain lengths. **(G)** Effect on the Y-to-Z ratio as a function of the step parameter $k_Y$ ($k_Z = 1$). **(H-I)** Analysis of cytokine blockade effects (cf. Figure 1, main text). **(H)** Effect on the ThX-to-ThY ratio in an RTM and SSM model as a function of the perturbation start time (duration of perturbation T=1d). Y-axis represents the log2-fold-change compared to a simulation with no perturbation. **(I)** Effect on the curve difference (same as in (H)) for variation of the step parameter k at early, intermediate and late perturbation times.

##### Figure S2: Supplementary data to Figure 2.

**(A-C)** Quality controls for kinetic transcriptome data (cf. Table 1 and Figure 2). **(A)** Normalized gene expression across different data sets. **(B)** Fitting error distribution (root-mean-square) for kinetic genes that were fitted using a response-time model or exponential model. **(C)** Number of kinetic genes categorized as “gamma”, “expo”, “bimodal” or “other” across data sets (cf. Figure 2B). **(D)** Estimated response-time distributions for proliferation and differentiation data (cf. Table 2). **(E)** Best-fit gamma distributions for the data shown in (D). **(F)** Example kinetics for genes in the “gamma” and “expo” categories. Panel titles indicate fit quality for the best-fit gamma-distribution and the closest Erlang distribution. **(G)** Example kinetics for genes in the bimodal and other categories. **(H)** Estimated mean and standard deviation for best-fit parameters $\alpha$, $\beta$ of individual effector cell differentiation modules (see Methods). **(I)** Normalized expression values for gene modules in different kinetic data sets (cf. Figure 2C).

##### Figure S3: Supplementary data to Figure 3.

**(A)** Cartoon model of the “Mixed” proliferation model with restimulation. At each restimulation event, either the clock of the timer model is reset, or IL-2 secretion is induced in effector cells, or both. **(B)** Model simulations with restimulation in an IL-2, Timer and Mixed model version. Each line represents a single simulation (cf. Supplementary Text). **(C)** Left: Model simulations with restimulation in the IL-2, Timer and Mixed Models for low IL-2 secretion of effector cells (10 molecules cell$^{-1}$ s$^{-1}$). Filled areas represent the maximum range of observed dynamics. Right: Distribution of cell numbers at day 60 for the simulations shown in the left panel. **(D)** Dependence of Response Size and Peak Time in the Mixed model for variation of IL-2 and Timer specific parameters. Effect size denotes log2 fold-change with respect to simulation results with minimal readout values (cf. Figure 3E).

##### Figure S4: Supplementary perturbation and sensitivity analysis.

**(A)** Sensitivity analysis at day 9 post acute (left panel) and chronic (right panel) infection, analogous to Figure 4H. **(B)** Th1 cell numbers in model simulations for early, intermediate and late perturbation times, analogous to Figure 5B.

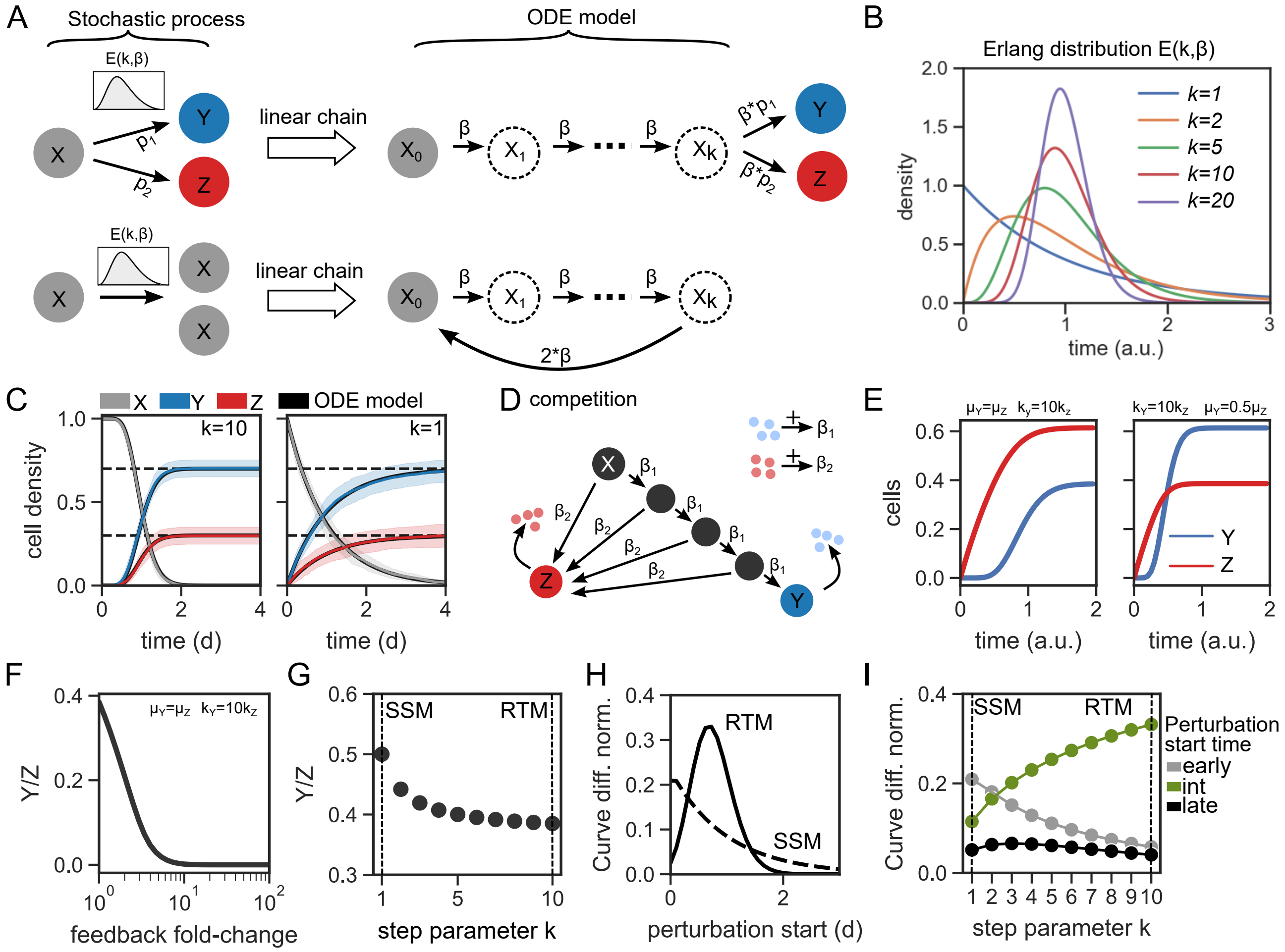


Burt et al. Figure S1

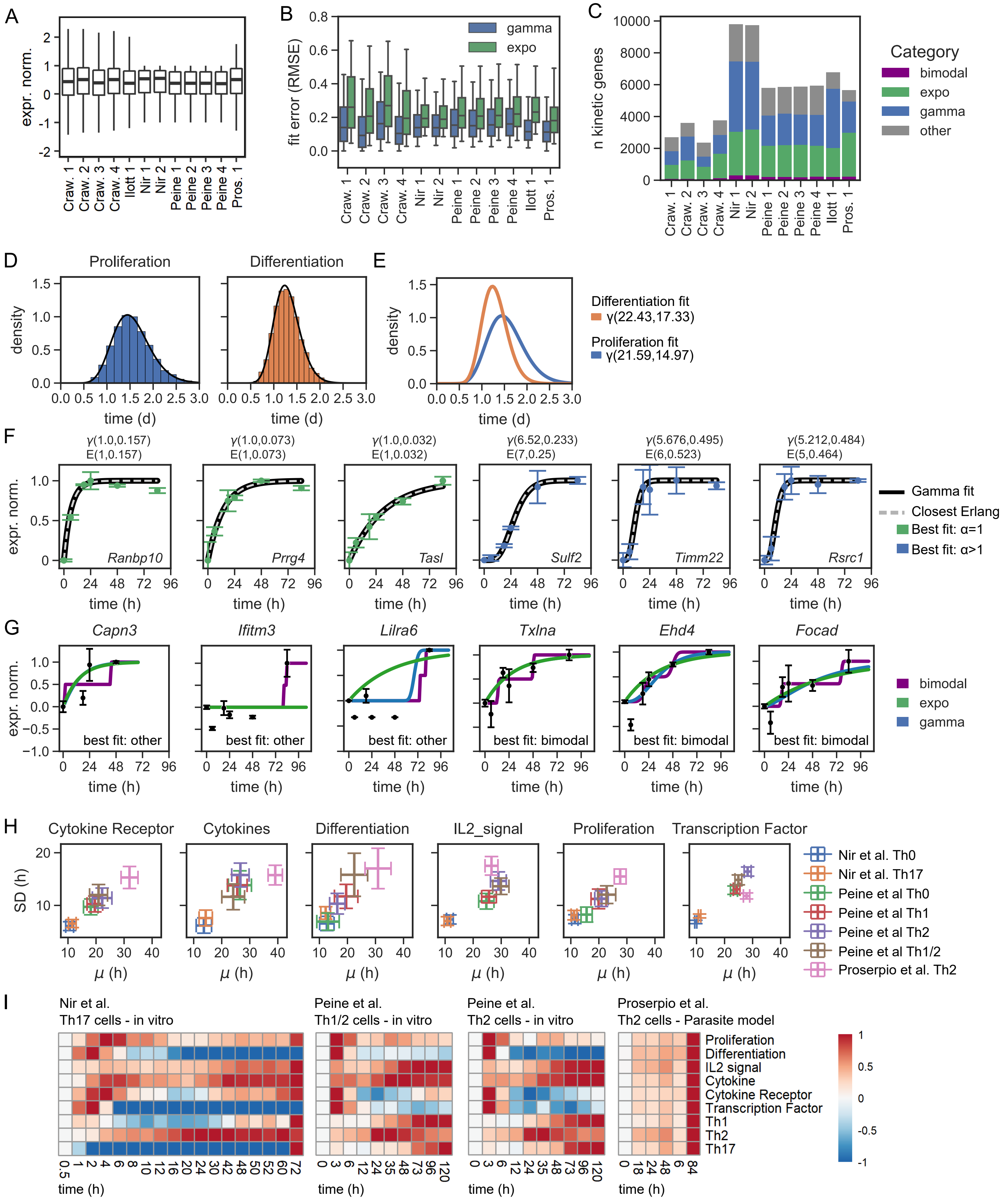


Burt et al. Figure S2

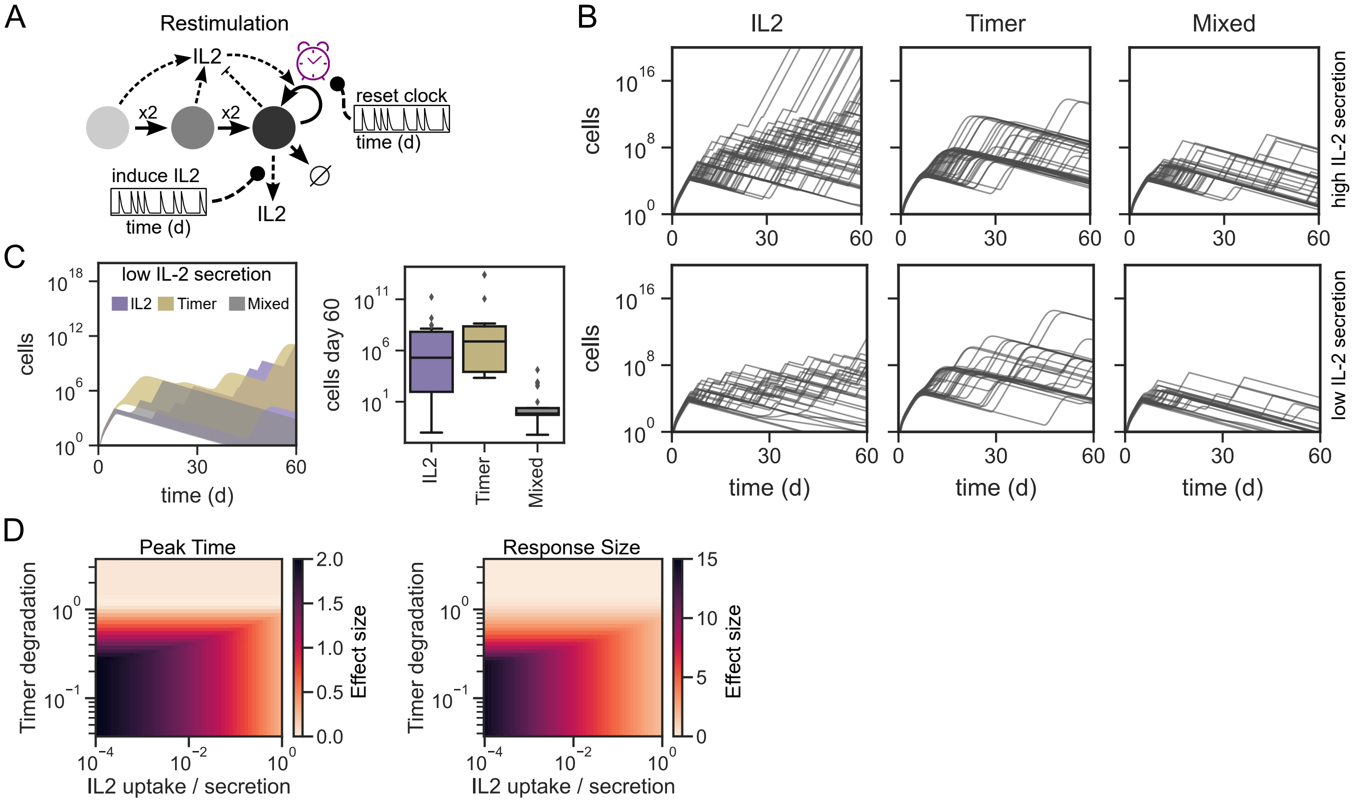


Burt et al. Figure S3

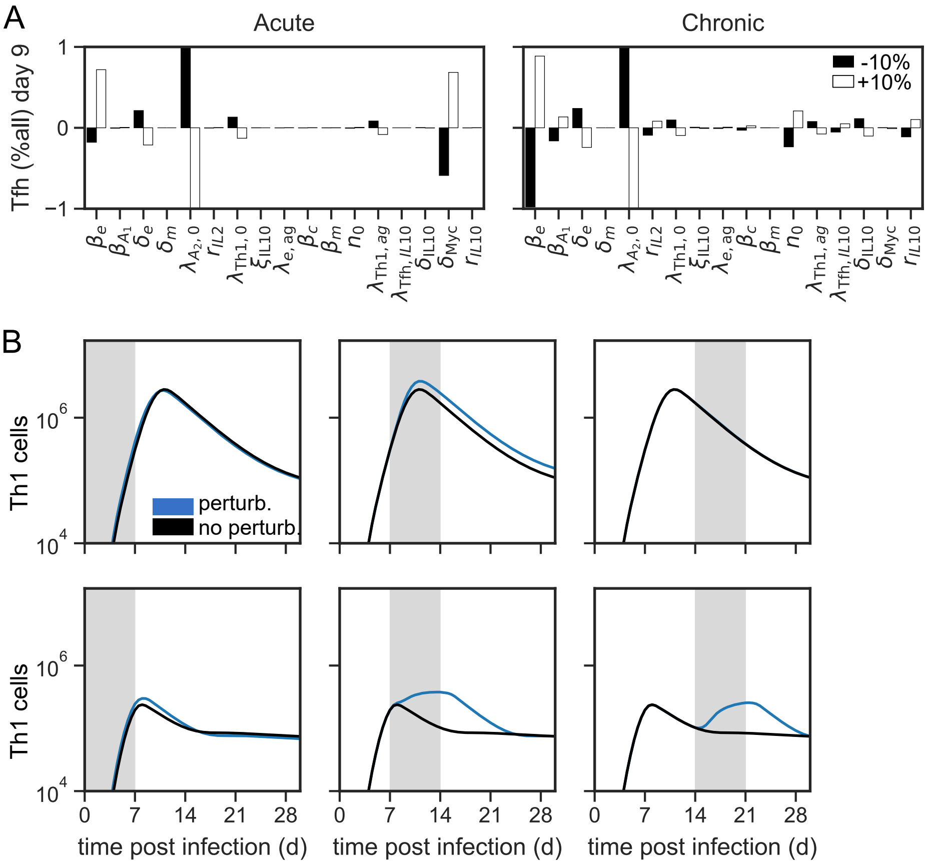


Burt et al. Figure S4
